# Elucidating relationships between *P.falciparum* prevalence and measures of genetic diversity with a combined genetic-epidemiological model of malaria

**DOI:** 10.1101/2020.08.27.269928

**Authors:** Jason A. Hendry, Dominic Kwiatkowski, Gil McVean

## Abstract

There is an abundance of malaria genetic data being collected from the field, yet using this data to understand features of regional epidemiology remains a challenge. A key issue is the lack of models that relate parasite genetic diversity to epidemiological parameters. Classical models in population genetics characterize changes in genetic diversity in relation to demographic parameters, but fail to account for the unique features of the malaria life cycle. In contrast, epidemiological models, such as the Ross-Macdonald model, capture malaria transmission dynamics but do not consider genetics. Here, we have developed an integrated model encompassing both parasite evolution and regional epidemiology. We achieve this by combining the Ross-Macdonald model with an intra-host continuous-time Moran model, thus explicitly representing the evolution of individual parasite genomes in a traditional epidemiological framework. Implemented as a stochastic simulation, we use the model to explore relationships between measures of parasite genetic diversity and parasite prevalence, a widely-used metric of transmission intensity. First, we explore how varying parasite prevalence influences genetic diversity at equilibrium. We find that multiple genetic diversity statistics are correlated with prevalence, but the strength of the relationships depends on whether variation in prevalence is driven by host- or vector-related factors. Next, we assess the responsiveness of a variety of statistics to malaria control interventions, finding that those related to mixed infections respond quickly (~ months) whereas other statistics, such as nucleotide diversity, may take decades to respond. These findings provide insights into the opportunities and challenges associated with using genetic data to monitor malaria epidemiology.

**Author summary:** Knowledge of how the prevalence of *P.falciparum* malaria varies, either between regions or through time, is critical to the operation of malaria control programs. Yet obtaining this information through traditional methods is fraught with challenges. Parasite genetic data is increasingly accessible, and may provide an alternative means to estimate *P.falciparum* prevalence in the field. However, our understanding of how the genetic diversity of parasite populations relates to prevalence is limited, and suitable models to guide our understanding are largely lacking. Here, we merge two classical models – the Ross-Macondald and the Moran – to produce a framework in which the relationships between parasite genetic diversity and prevalence can be explored. We find that several genetic diversity statistics are correlated with prevalence, although to differing degrees, and over different time scales. Overall, statistics related to mixed infection are robustly and rapidly responsive to changes in prevalence, suggesting they may be a useful focal point for the development of malaria surveillance methods that harness genetic data.

## Introduction

It is widely accepted that relationships exist between the regional epidemiology of malaria and the genetic diversity of local parasite populations. For example, the evolution of antimalarial drug resistance, patterns of parasite migration and variation in transmission intensity may all have relationships with population genetic diversity (reviewed in [1–7]). In most cases, however, the precise nature of these relationships remains unclear. From a modelling perspective, exploring these relationships would require that both genetic processes (including mutation, drift and meiosis) and epidemiological ones (including the transmission dynamics and life cycle of malaria) are combined into a single framework. At present such integrated models are rare, yet without them, parasite genetic data will be under-utilized as a resource for malaria surveillance.

One epidemiological parameter of central importance to malaria surveillance is transmission intensity, as it is used by National Malaria Control Programs (NMCPs) to prescribe malaria control interventions and assess their efficacy [8]. NMCPs can attempt to measure transmission intensity in a variety of ways, including with the basic reproduction number (*R*_0_), the entomological inoculation rate (*EIR*), parasite prevalence (*P R* or *P f P R* if the focus is *P. falciparum*), or rates of clinical incidence (reviewed in [9]). However, there are well-documented issues with all of these measurement approaches. Though a theoretical gold-standard, *R*_0_ is difficult to measure in practice, with estimation methods relying either on exploiting equilibrium relationships to other measures of transmission, or formulae involving several poorly-characterised parameters [9, 10]. The *EIR* suffers from small-scale variability in mosquito density, a lack of standardisation across mosquito catching methods, and difficulties associated with catching sufficient mosquitoes when transmission intensities are low [9, 11, 12]. Rates of clinical incidence are confounded by variation in acquired immunity and treatment seeking behaviour, as well as incomplete record keeping [12]. Parasite prevalence is the most widely collected measure and has been used as the basis of large-scale maps [13–15], yet it requires prohibitively extensive sampling at low transmission intensities [12], and must address biases in detection power that may arise from infection-course and age-dependent variation in parasitemia [9, 16]. Thus, a means to either estimate or improve existing estimates of transmission intensity with genetic data would be valuable.

The problem of estimating an epidemiological parameter like transmission intensity from genetic data is superficially similar to the demographic inference problems that are commonly encountered in population genetics. For example, a multitude of methods now exist for estimating effective population size (*N*_*e*_) from genetic data [17–20], and it could be hypothesized that the regional transmission intensity of malaria is a function of the *N*_*e*_ of the local parasite population. However, there are at least two challenges unique to epidemiological inference from malaria genetic data that make it a distinctive and more difficult problem.

First, it is not clear that the classical models in population genetics (including the Wright-Fisher, Moran, and other models that converge to Kingman’s *n*–Coalescent in the ancestral limit [21]) that are often employed by demographic inference methods are suitable for malaria. The life cycle of *P. falciparum* involves oscillating between human host and mosquito vector populations, which may be of different sizes, and may also induce different rates of drift and mutation. Within both the host and the vector, parasite populations likely experience bottlenecks (for example, as the ookinetes penetrate the midgut wall of the vector), exponential growth phases (merozoites replicating in the blood of the host), and interactions with the host or vector immune system. Indeed, there has been work demonstrating that the malaria life cycle simultaneously intensifies drift and selection; a result contrary to what is expected under a Wright-Fisher model [22]. So, while classical models in population genetics have the advantage of being extensively studied and mathematically tractable, with a known set of relationships between equilibrium genetic diversity statistics and demographic parameters, they do not readily apply to *P. falciparum*. Finally, even if these models did apply, the relationships between demographic parameters (such as *N*_*e*_) and epidemiological ones (such as transmission intensity) would need to be defined.

Conversely, the epidemiological models that have been designed to reflect malaria biology and transmission do not explicitly incorporate genetic processes. The most well known class of epidemiological models are the so-called “compartment-based” models, where individual hosts and vectors transition between compartments which can represent a variety of disease states (such as susceptible, infected, or immune), and the overall population is represented by the total number of hosts and vectors occupying each state (reviewed in [23, 24]). A canonical compartment model for malaria is the Ross-Macdonald, where hosts and vectors can be either susceptible or infected [24]. Conveniently, in these models, the equilibrium prevalence of infected hosts and vectors is a function of the parameters specifying the transition rates between compartments. However, the absence of genetic processes (and indeed, individual parasites) means they offer no insight into how parasite genetic diversity relates to these transition rates or, as a corollary, any epidemiological parameters derived from them. Thus, at least with respect to malaria, the traditional modelling landscape cannot address questions that involve both genetic data and epidemiology.

A second challenge specific to epidemiological inference using genetic data is that the ultimate aim is often to inform disease control and, as a result, the time-dimension of the inference is of critical importance. Many demographic inference methods base their estimates of *N*_*e*_ on distributions of coalescent times between segments of DNA. As these coalescent events occur on average *N*_*e*_ generations in past, the estimates are historical; reflecting the average population size over hundreds or thousands of generations. Such approaches are not suitable for disease control, where policy decisions need to be made on the basis of information about the near-present, or predictions about the future.

An integrated genetic-epidemiological model was developed previously to analyse *P. falciparum* single-nucleotide polymorphism (SNP) data, collected during a period of intensified intervention in Thi`es, Senegal [25]. Fit only to 24-SNP barcodes, the model independently corroborated a decline and rebound in transmission intensity (measured by *R*_0_ ), thus demonstrating the potential utility of genetic data for malaria surveillance [25]. However, as the availability of *P. falciparum* whole-genome sequencing (WGS) data has since increased, there is a need for modelling frameworks that can investigate the broader suite of genetic diversity statistics calculable from WGS data. Moreover, the model developed by *Daniels et al.* [25] was tailored for a specific application, and there are additional features of malaria epidemiology that remain to be investigated. These include the relevance of the vector population, the effects of intra-host and -vector evolution, and the influence of super-infection and co-transmission on genetic diversity.

Here, we aim to address the lack of integrative genetic-epidemiological models by developing a new forward-time model called forward-dream. forward-dream merges the Ross-Macdonald model with a continuous-time intra-host and intra-vector Moran model, and further incorporates meiosis within the vector (allowing for multiple oocysts), multiple infection (by either super- or co-infection), and a representation of the transmission bottlenecks. Implemented as a stochastic simulation, we use the model to explore relationships between measures of genetic diversity and parasite prevalence, both at equilibrium and in response to malaria control interventions that perturb equilibrium. We confirm that a variety of genetic diversity statistics are correlated with parasite prevalence, although to varying degrees and over different time-scales. In addition, we find that interventions that affect the duration of infection in hosts have a greater influence on parasite genetic diversity than those that influence vector biting rate or density. Overall, our results suggest that statistics based on the complexity of infection (C.O.I.) are strongly, robustly, and rapidly responsive to changes in prevalence, highlighting their potential value for malaria surveillance.

## Results

### Developing a model of *P.falciparum* malaria transmission and evolution

We developed an agent-based simulation of *P.falciparum* malaria incorporating features of its transmission and life cycle, as well as explicitly modelling the genetic material of parasites. Our integrated genetic-epidemiological model is called forward-dream (**forward**-time **d**rift, **r**ecolonisation, **e**xtinction, **a**dmixture and **m**eiosis) and is comprised of three layers: (1) an epidemiological layer, which controls how malaria spreads through a population of hosts and vectors and reaches equilibrium; (2) an infection layer, which controls the behaviour of malaria parasites during individual transmission events and within individual hosts and vectors; and (3) a genetic layer, which controls how the genetic material of individual parasites is represented, mutated and recombined. We describe each layer below and provide additional information, including a discussion of parameterisation, in the supplementary materials.

#### Epidemiological layer

For the epidemiological layer we implemented the Ross-Macdonald model (reviewed in [24]), in which a fixed number of hosts (*N*_*h*_) and vectors (*N*_*v*_) alternate between susceptible and infected based on four fixed rate parameters (Fig 1a). The model can be described by two coupled differential equations:

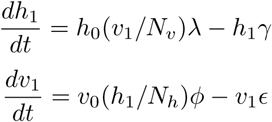

where *h*_0_ and *h*_1_ are the number of susceptible and infected hosts, respectively, with *N*_*h*_ = *h*_0_ + *h*_1_; and *v*_0_ and *v*_1_ are the number of susceptible and infected vectors, respectively, with *N*_*v*_ = *v*_0_ + *v*_1_. Note that in this model *λ* and *φ* are compound parameters representing the rate at which hosts and vectors become infected. In particular,

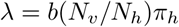

where *b* is the daily vector biting rate, (*N*_*v*_/*N*_*h*_) gives the vector density, and *π*_*h*_ is the probability that an infectious bite from a vector produces an infected host (the vector-to-host transmission efficiency). Similarly, we have,

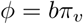

where *b* is again the daily vector biting rate and *π*_*v*_ is the probability that a vector that bites an infected host becomes infected (the host-to-vector transmission efficiency). We allow for mixed infections however, for the epidemiological layer, they have no consequence: both clonal and mixed infections are assigned to the infected compartments (*h*_1_ or *v*_1_).

**Figure 1.**
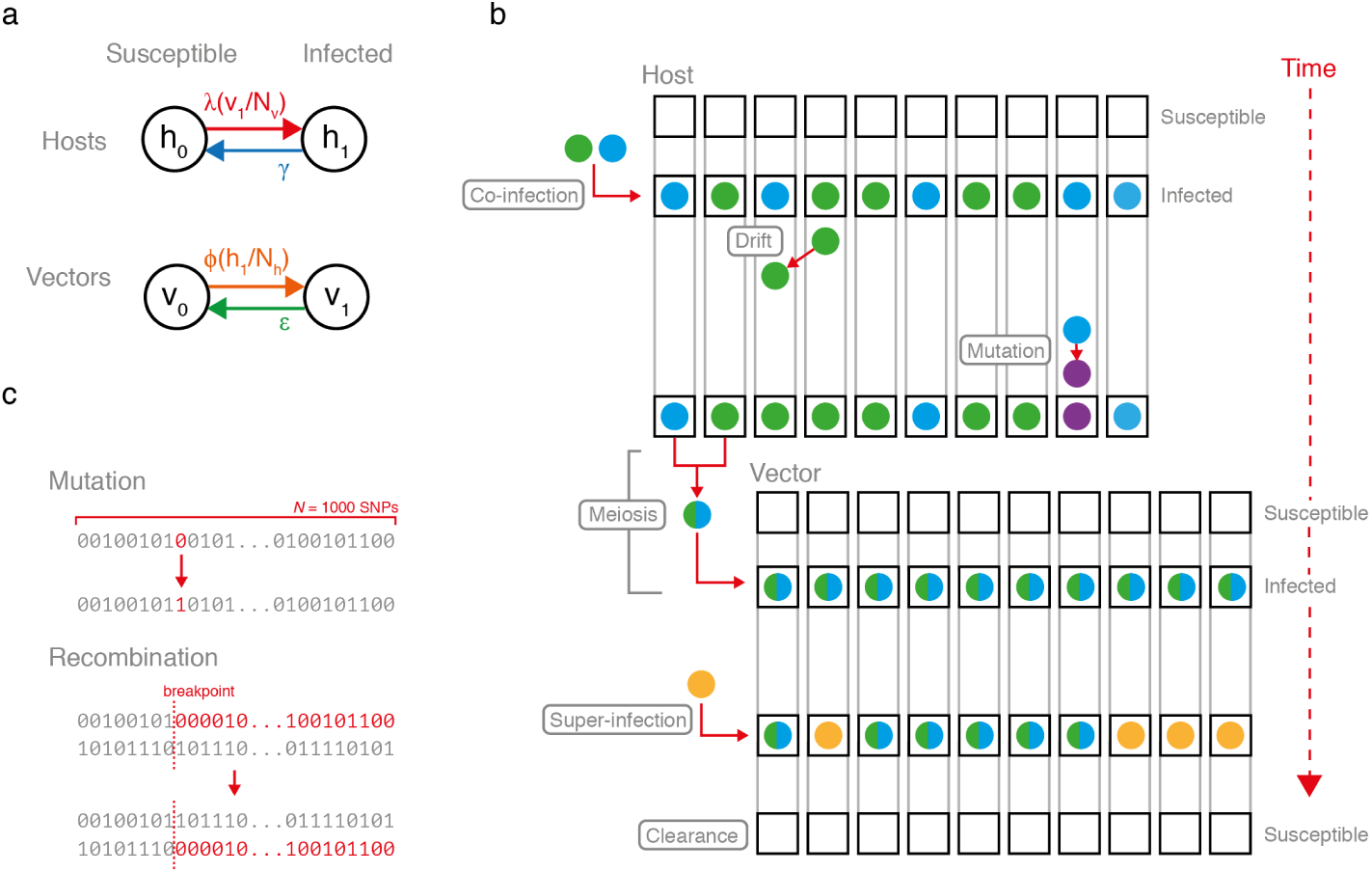
Schematic of forward-dream. (a) The epidemiological layer. Hosts and vectors oscillate between susceptible and infected compartments according to a Ross-Macdonald model. (b) The infection layer. The capacity of individual hosts and vectors to be infected is represented by a fixed number of sub-compartments (black boxes) each which can harbour a unique parasite genome (colored circles). In a susceptible host/vector, all sub-compartments are empty. Upon infection, all sub-compartments are populated. Drift and mutation occur among sub-compartments according to a continuous-time Moran model. Parasite genomes undergo meiosis during transmission from host to vector. Super-infection can occur, resulting in an average of half of all sub-compartments being replaced with newly transmitted parasite genomes. Note that the infection layer can be nested within the Ross-Macdonald model. (c) The genetic layer. The genome is represented by a fixed-length array of 0’s and 1’s. Mutation is reversible, converting 0 to 1 or 1 to 0. Recombination occurs during meiosis.

Note that the epidemiological layer dictates the equilibrium prevalence of infection in hosts (*X*_*h*_ = *h*_1_/*N*_*h*_) and vectors (*X*_*v*_ = *v*_1_/*N*_*v*_). In particular, the equilibrium prevalence in hosts is given by:

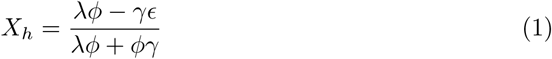

and in vectors by: 

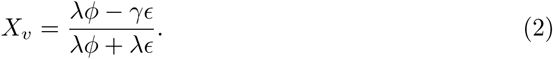

We have confirmed that our implementation of the Ross-Macdonald model in forward-dream converges to the expected equilibrium prevalence values (Fig 2a and S1 Fig). In total, the behaviour of the epidemiological layer is specified by seven parameters.

**Figure 2.**
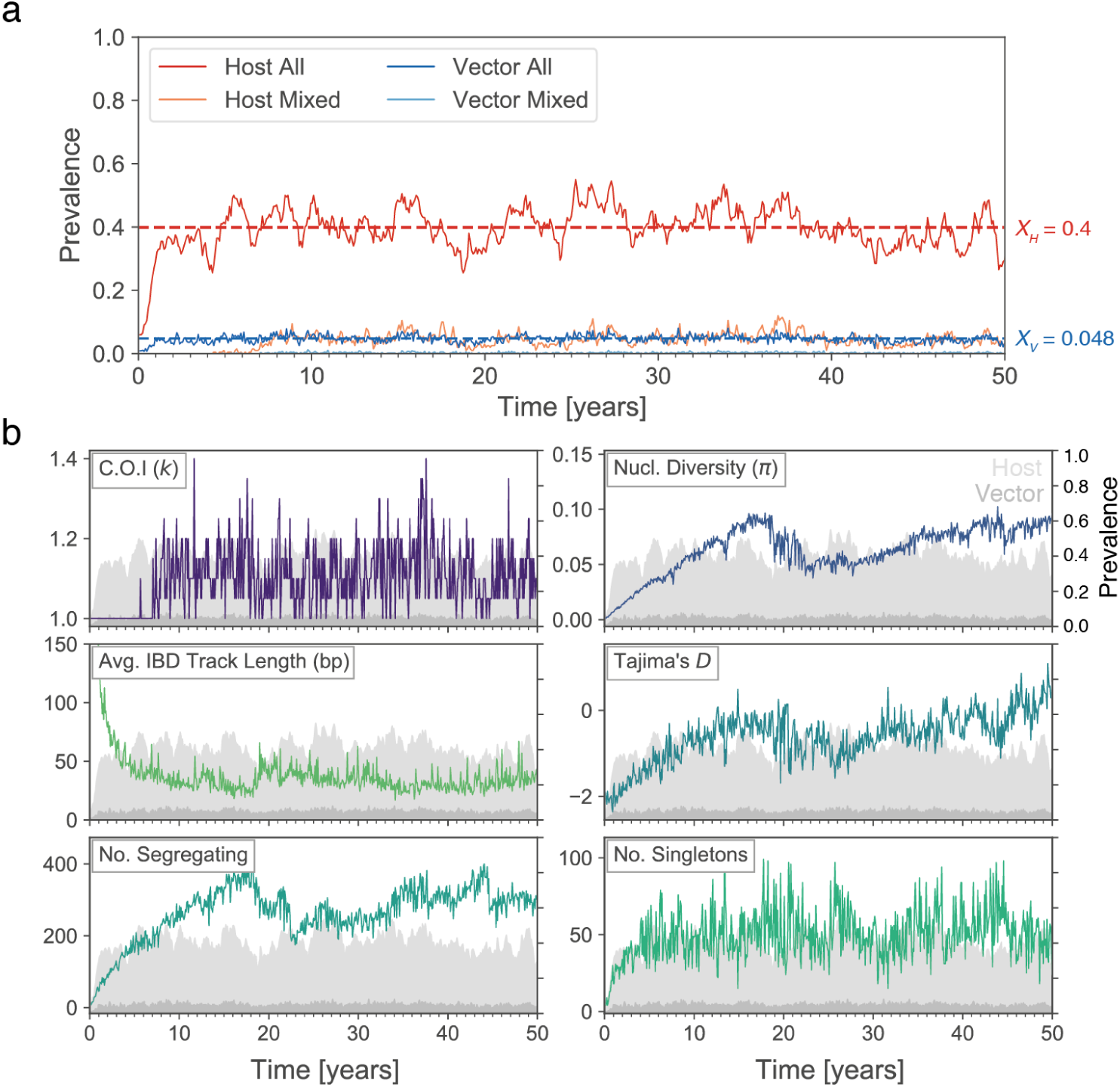
Simultaneously monitoring parasite prevalence and genetic diversity using forward-dream. (a) A forward-dream simulation seeded with ten infected hosts harbouring identical parasite genomes and run for 50 years. Prevalence of all infected hosts (’Host All’, which corresponds to *P f P R*) and multiply-infected hosts (’Host Mixed’) is indicated by red and pink lines, respectively. The same is shown for vectors in blues. The prevalence of infected hosts and vectors fluctuates around their Ross-Macdonald equilibrium values (*X*_*h*_ and *X*_*v*_), indicated with red and blue horizontal lines, respectively. (b) The same simulation as in (a), but visualizing six genetic diversity statistics that were computed by collecting parasite genomes from twenty randomly selected infected hosts every 30 days. The statistics are defined in S1 Table. For reference, the light and dark grey shaded areas show the host and vector prevalence (right y-axis), corresponding to the red and dark blue lines in panel (a).

#### Infection layer

The infection layer specifies the biology of individual infection processes and transmission events. This includes a representation of: (i) the infection capacity of hosts and vectors, (ii) the evolution of infections within hosts and vectors, and (iii) the transmission of infection between hosts and vectors.

We model an individual host’s capacity to be infected with *n*_*h*_ within-host sub-compartments (Fig 1b). When an individual host is in the susceptible state (i.e. is uninfected), all *n*_*h*_ sub-compartments are empty. When an individual host becomes infected, all *n*_*h*_ sub-compartments simultaneously become populated. Each sub-compartment can potentially harbour a unique parasite genome (such that the maximum complexity of infection *k* = *n*_*h*_), although typically multiple sub-compartments will be occupied by identical, or near-identical, genomes. For example, if a host is infected by a vector carrying a single distinct parasite genome, all *n*_*h*_ sub-compartments will initially be occupied by that genome. Alternatively, if a host is co-infected by a vector carrying two distinct parasite genomes, its sub-compartments will be a mixture of the two genomes.

Moving forward in time, the *P. falciparum* infection of a host evolves according to a continuous-time Moran process for a population with *n*_*h*_ individuals, parameterised by a drift rate (*d*_*h*_) and mutation rate (*θ*_*h*_) [26]. We have confirmed that our implementation of the Moran process yields fixation times consistent with theoretical expectation (S2 Fig). The Moran process continues until the infection is cleared and the host returns to the susceptible state, with all sub-compartments becoming simultaneously empty. For hosts, clearance occurs at rate *γ*, as specified in the epidemiological layer. The infection of a vector evolves according to a comparable process as in the host, but with a set of parameters *n*_*v*_, *d*_*v*_, *θ*_*v*_, and *ϵ*.

In forward-dream, a *P. falciparum* infection is transmitted from a vector to a host through a transmission bottleneck, such that not all of the parasites within the infectious vector establish themselves in the host. In particular, a random subset of all parasites 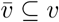 (where 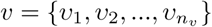 is the set of all parasites in the vector) is transmitted. The number of transmitted parasites 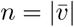 is drawn from a truncated binomial:

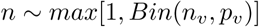

This results in an average of 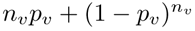 parasites passing through the transmission bottleneck; the size of the bottleneck is controlled by *p*_*v*_. For all of the simulations presented here, *p*_*v*_ = 0.2 and *n*_*v*_ = 10, resulting in an average of ≃ 2 parasites passing through the bottleneck.

Note that in cases where *n* > 1 and 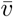 contains unique parasites, co-infection may occur. If the vector is infecting a susceptible (i.e. uninfected) host, *n*_*h*_ parasites are drawn with replacement from 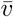 with each having an equal probability (1*/n*) of being drawn. These *n*_*h*_ parasites then populate the *n*_*h*_ host sub-compartments. Alternatively, super-infection occurs if the host is already infected. Defining the parasites already within the host by the set 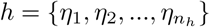, we first create the union 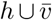. From this set *n*_*h*_ parasites are drawn, where each parasite has probability 1/2*n*_*h*_ or 1/2*n* of being drawn, if it is from *h* or 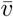, respectively. As a consequence, on average super-infection results in half of the within-host compartments being occupied by new parasites.

The transmission from host to vector is the same as above, but with the addition of meiosis. In brief, the *n* parasite strains selected at random from the infected host may undergo meiosis before populating the the *n*_*v*_ within-vector sub-compartments. The meiosis model is based on a simplified implementation of our previously published meiosis simulator, pf-meiosis [27]. It includes multiple oocysts, allowing for parallel rounds of meiosis to occur during a single transmission event, with the number of oocysts being drawn from a truncated geometric: ~*min*[1, *Geo*(*p*_*oocysts*_)].

In total, the infection layer is specified by nine parameters.

#### Genetic layer

The genetic layer of forward-dream describes the malaria genome model. We represent the genetic material of an individual parasite as a single fixed-length array of zeros and ones, defined by the parameter *N*_*snps*_ (Fig 1c). In effect, this array represents a single chromosome marked with *N*_*snps*_ single-nucleotide polymorphisms (SNPs). Mutation is reversible and the recombination rate is constant and scaled with respect to *N*_*snps*_, such that an average of one cross-over event occurs per bivalent during meiosis. As a result the only parameter specific to the genome evolution layer is *N*_*snps*_.

Overall, forward-dream is specified by 17 parameters (Table 1). It is implemented in Python and available on GitHub at https://github.com/JasonAHendry/fwd-dream.

**Table 1.** Complete list of simulation parameters for forward-dream. Values given represent those of a simulation with a host (*X*_*h*_) and vector (*X*_*v*_) prevalence of 0.65 and 0.075, respectively. This corresponds to the Initialise epoch of all malaria control intervention simulations (see below). For details on how parameter values were selected see the Supporting Materials.

| Parameter | Definition | Value |
| --- | --- | --- |
| $N_h$ | Number of hosts | 400 |
| $N_v$ | Number of vectors | 2000 |
| $b$ | Biting rate (per vector per day) | 0.25 |
| $\pi_h$ | Vector-to-host transmission efficiency | 0.1 |
| $\pi_v$ | Host-to-vector transmission efficiency | 0.1 |
| $\gamma$ | Host infection clearance rate | 0.005 |
| $\epsilon$ | Vector infection clearance rate | 0.2 |
| $n_h$ | Maximum C.O.I. ($k$) for hosts | 10 |
| $n_v$ | Maximum C.O.I. ($k$) for vectors | 10 |
| $d_h$ | Drift rate in hosts (events per day per host) | 1 |
| $d_v$ | Drift rate in vectors (events per day per vector) | 1 |
| $\theta_h$ | Probability of mutation event given drift has occurred in hosts | 0.0001 |
| $\theta_v$ | Probability of mutation event given drift has occurred in vectors | 0.0001 |
| $p_h$ | Probability a given parasite genome passes through host-to-vector transmission bottleneck | 0.2 |
| $p_v$ | Probability a given parasite genome passes through vector-to-host transmission bottleneck | 0.2 |
| $p_{\text{oocysts}}$ | Number of oocysts drawn from $\sim \text{Geo}(p_{\text{oocysts}})$ | 0.5 |
| $N_{\text{snps}}$ | Number of SNPs modelled per parasite genome | 1000 |

### Relationships between parasite prevalence and genetic diversity at equilibrium

Within the forward-dream framework, it is straightforward to monitor the fraction of hosts that are infected (*h*_1_/*N*_*H*_), which corresponds to the most ubiquitously collected measure of transmission intensity, parasite prevalence (*P f P R*) (Fig 2a). It is also possible to monitor the fraction of vectors infected (*v*_1_/*N*_*V*_), which corresponds closely to the sporozoite rate (*SP*). Finally, the genetic diversity of the parasite population can be monitored by collecting parasite genomes from infected hosts and simulating DNA sequencing (see Materials and Methods) (Fig 2b).

We sought to use forward-dream to elucidate relationships between *P f P R* and the genetic diversity of the parasite population. To this end, we varied *P f P R* across simulations and observed the resulting differences in parasite genetic diversity. Within the Ross-Macdonald framework, *P f P R* is a function of the four rate parameters (see Eq. 1). In nature, what underlies prevalence differences observed between two geographies or points in time is often unknown, and likely the outcome of a myriad of epidemiological and environmental factors. To achieve different *P f P R* values in forward-dream, we choose three parameters to vary separately: (i) the human infection clearance rate (*γ*), which may vary between sites if, at one site there is quicker recourse to treatment, or differing proportions of symptomatic and asymptomatic individuals; (ii) the vector biting rate (*λ*), which may vary as a consequence of differences in vector species, environmental conditions, or the presence of bednets; and (iii) the number of vectors (*n*_*v*_) – which is influenced by climate and weather, local geography, and also by insecticide-based interventions (see [28]). We tuned each of these parameters to achieve equilibrium *P f P R* values that varied from 0.2 to 0.8 in a population of four-hundred human hosts (S3 Fig). We note that prevalence values less than 0.2 are of interest, however, they produce frequent stochastic extinction in forward-dream (especially over long time periods) and thus were not studied here. At each prevalence value, forward-dream was seeded with ten infected hosts carrying identical parasites and then run to equilibrium. After reaching equilibrium, simulations were continued for an additional 10 years, during which time parasite genomes were collected every thirty days from twenty infected hosts selected at random. From these collected parasite genomes we could track the behaviour of a suite of genetic diversity statistics through time and construct their distributions at a given fixed prevalence.

The genetic diversity statistics calculated are described in S1 Table. The statistics can be divided into three broad categories: (i) those related to mixed infections, which includes the fraction of mixed samples and the mean complexity of infection (C.O.I.); (ii) those that are related to the size and shape of the genetic genealogy of the sample, which includes the number of segregating sites, the number of singletons, nucleotide diversity (*π*), Watterson’s Theta (*θ*_*w*_), and Tajima’s *D*; and (iii) those that summarize the structure of identity-by-descent (IBD) within the population, including, between a pair of samples, the average fraction of the genome in IBD, the average number of of IBD tracks, and the average length of an IBD track. We note that these statistics are not independent, indeed many are co-linear (S7 Fig), however they reflect commonly-used measures of genetic diversity in population genetics.

We plotted the distributions of each of these ten statistics across different *P f P R* values, partitioned by which epidemiological parameter was varied (Fig 3a and S4 Fig to S6 Fig). We used linear regression to determine the fraction of the variance (*r*^2^) in each genetic diversity statistic that could be explained by variation in *P f P R*. A summary of the results of this analysis are shown in Fig 3b. Of the ten genetic diversity statistics, all but Tajima’s *D* and the number of singletons have a substantial proportion of their variance (> 20%) explained by *P f P R*, regardless of the underlying epidemiological cause. All of the genetic diversity statistics have a positive relationship with prevalence, except for the average fraction of IBD and average IBD track length, which decrease in higher prevalence regimes.

**Figure 3.**
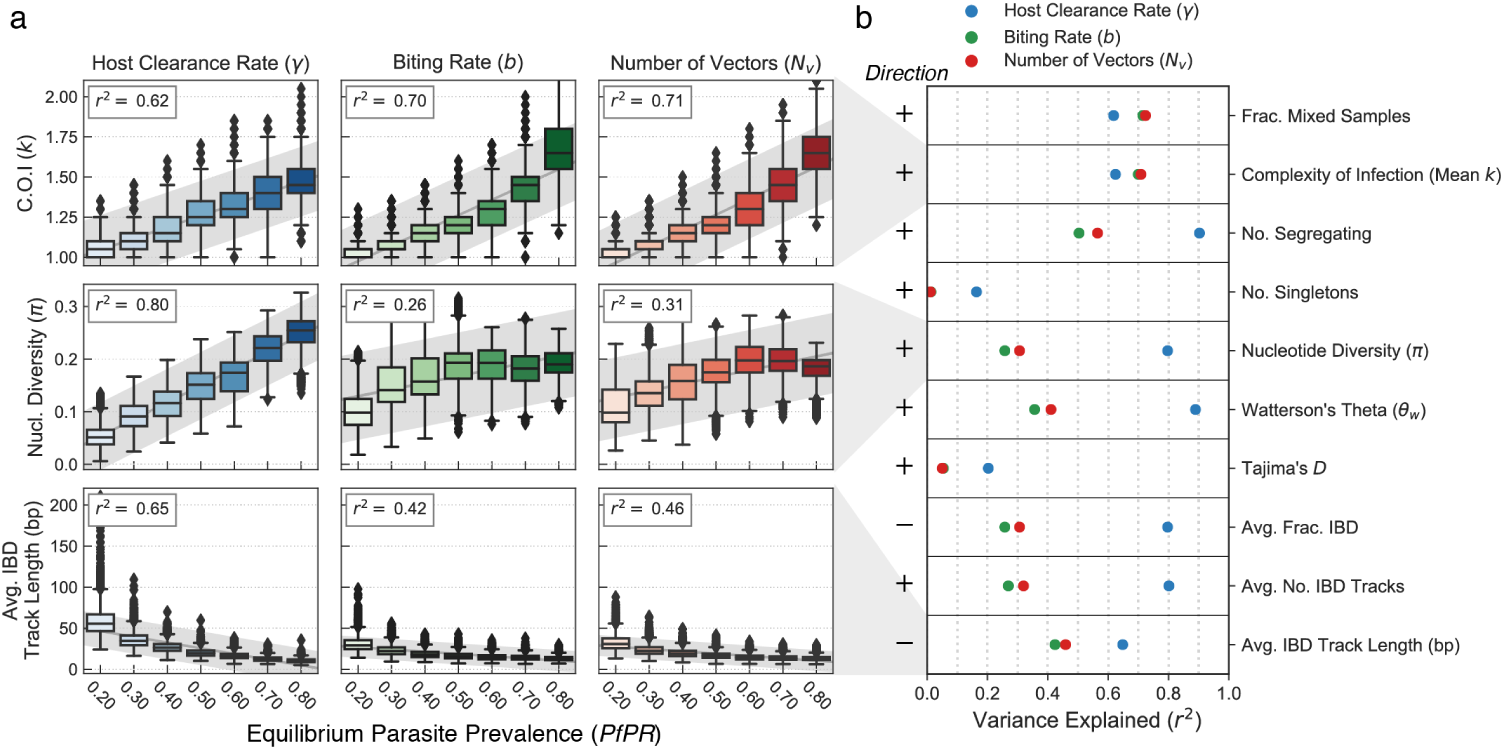
Equilibrium relationships between genetic diversity and parasite prevalence. (a) Boxplots showing the equilibrium parasite prevalence in hosts (x-axis) versus genetic diversity of the parasite population, by different statistics. Each individual box summarizes 30 replicate simulations at the indicated prevalence; in each, 20 infected hosts were sampled every 30 days for ten years to collect parasite genomes from which diversity statistics were computed. Left, middle, and right columns show relationships when equilibrium parasite prevalence is varied as a function of the host clearance rate (*γ*, in blue), vector biting rate (*b*, in green) or number of vectors (*N*_*v*_, in red). The variance explained in an ordinary linear regression (*r*^2^) is shown at the top left for each relationship, and in (b) the variance explained is shown across a larger panel of genetic diversity statistics. Slope of the line of best fit is indicated at left (+, increasing; -, decreasing). Boxplots for all statistics in (b) can be found in the supplementary figures.

Nevertheless, there are pronounced differences in the variance explained by *P f P R* when different epidemiological parameters drive variation in prevalence. In particular, varying host clearance rate (*γ*) results in a significantly higher variance explained for all of the genealogy and IBD statistics. For example, the variance in nucleotide diversity explained by *P f P R* drops from 80% to 26% when parasite prevalence is modulated by the vector biting rate (*b*), instead of the host clearance rate (*γ*). In contrast, the two statistics related to mixed infections have high *r*^2^ values (> 60%) regardless of which of the epidemiological parameters underlies the parasite prevalence change.

### Non-equilibrium relationships between parasite prevalence and genetic diversity

An important application of our model is in settings where malaria control interventions are actively occurring. In such cases, and also in cases with seasonal variation in parasite prevalence, it is unlikely that the parasite population will be in equilibrium. Thus, our second aim with forward-dream was to explore which measures of genetic variation are most predictive of instantaneous *P f P R* in non-equilibrium settings.

#### Malaria control interventions

In order to understand how genetic diversity statistics relate to *P f P R* in scenarios where a malaria control intervention has been deployed, we developed a framework where individual forward-dream simulations pass through three distinct epochs: Initialise, Crash and Recovery (Fig 4). In the Initialise epoch, simulations are run until equilibrium at a parasite prevalence of 0.65, under the parameters listed in Table 1. At the start of the Crash epoch, one of either the host clearance rate (*γ*), the vector biting rate (*b*), or the number of vectors (*N*_*v*_), is changed such that the new equilibrium *P f P R* is 0.2. The parameter change occurs incrementally over a period of thirty days following a logistic transition function, as to mimic the staged introduction of a malaria control intervention. As a consequence, the simulation leaves equilibrium, with host and vector prevalence declining. The Crash epoch is allowed to continue until the population has regained equilibrium. At the start of the Recovery epoch, we return the changed simulation parameter to its original value, again over thirty days following a logistic transition function. Again, this results in the simulation leaving equilibrium, with the *P f P R* increasing back to 0.65. As with the Crash epoch, the Recovery epoch continues until the population regains equilibrium. In summary, the three epoch model allows us to explore both a parasite population decline and rebound.

**Figure 4.**
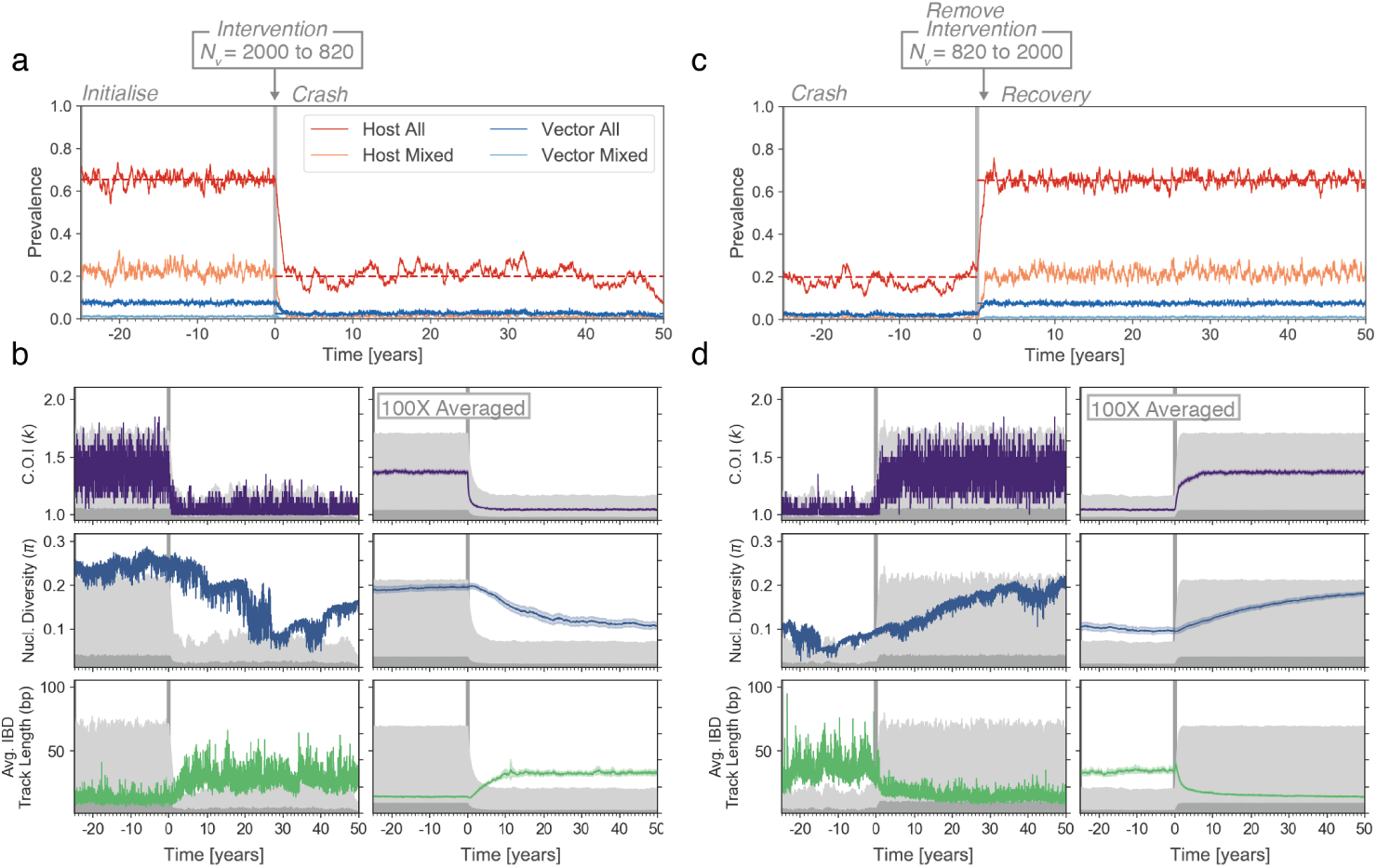
Responses of genetic diversity statistics to a crash and recovery in parasite prevalence. (a) An individual forward-dream simulation where the number of vectors (*N*_*v*_) is reduced at time zero (x-axis, indicated by grey vertical bar), resulting in a decline of parasite prevalence in hosts from 0.65 to 0.2. (b) Left column, the same simulation as in (a), but showing the response of three genetic diversity statistics (colored lines) to the prevalence change. Light and dark grey areas show host and vector prevalence, as in Fig 2. Right column, mean value of each genetic diversity statistics across 100 independent replicate simulations, with shading showing the 95% confidence interval. Average trajectories of additional statistics can be found in the supplementary materials. (c) Same simulation as in (a) at a later time, where the number of vectors is returned to its original value. Parasite prevalence increases back to 0.65. (d) Same as (b), but corresponding to the recovery shown in (c).

Throughout the three epochs parasite genomes are collected every five days from twenty hosts selected at random, allowing the same suite of genetic diversity statistics discussed above to be followed through time. An immediate observation was that in individual simulations, the trajectories of genetic diversity statistics were very “noisy”, tending to fluctuate considerably through time even in the absence of any change to simulation parameters (Fig 4 and S8 Fig to S10 Fig). Several compounding factors likely contribute to these fluctuations, including the stochastic nature of the epidemiological process (host and vector prevalence fluctuate through time around their equilibrium values), the stochastic nature of the genetic process (drift, mutation, and meiosis all occur randomly) and the variance introduced by sampling only 20 of 400 hosts.

To better discern the average behaviour of different statistics, we sought to smooth their trajectories by computing a rolling mean. However, we found that the timescale of random fluctuations differs greatly among genetic diversity statistics. Statistics such as the mean C.O.I. and the fraction of mixed samples fluctuate rapidly, and variation around their true mean could be substantially reduced (to <20% the original variance) by averaging the genetic data collected over roughly a month (S11 Fig). Yet we found that other statistics, such as nucleotide diversity, are more inertial in nature; they can trend up or down over very long time periods without any change to the underlying simulation parameters. Reducing their initial variance to an equivalent degree required averaging over 10 years worth of genetic data (S11 Fig).

Thus, we took an alternate approach to extract underlying trends: we averaged the trajectories of each statistic across 100 independent, replicate forward-dream simulations (Fig 4b, d and S12 Fig to S14 Fig). Several observations emerged. First and consistent with population genetic theory, Tajima’s *D* increases in the period where *P f P R* is declining (indicative of a contracting *N*_*e*_), and falls below zero in the period where *P f P R* is climbing (indicative of an expanding *N*_*e*_) [29] (S15 Fig). Second, there are marked differences in the rate at which different genetic diversity statistics respond to changing prevalence. For example, the mean C.O.I. responds faster than the average IBD track length, which in turn responds faster than nucleotide diversity. Finally, we noted that the rate at which a given genetic diversity statistics responds to a decline in *P f P R* may be different from the rate at which it responds to an increase in *P f P R* (Fig 4).

We developed two metrics to summarize the temporal responses of different genetic diversity statistics to changes in parasite prevalence in our simulations (Fig 5, see also Materials and Methods). Within the three epoch simulation framework, we could construct equilibrium distributions for each statistic before and after each prevalence change (i.e. equilibrium distributions for Initialise, Crash, and Recovery). Using these distributions, we computed a “detection time” (*t*_*d*_), which we define as the amount of time, following an intervention, until a given genetic diversity statistic takes on values outside of its pre-intervention equilibrium. In effect, *t*_*d*_ is an estimate of how long it takes to detect that a change in parasite prevalence has occurred by monitoring a given genetic diversity statistic through time. We also computed an “equilibrium time” (*t*_*e*_), which we define as the amount of time until a given genetic diversity statistic reaches its new, post-intervention equilibrium. Note that these metrics were designed to be informative summaries of the simulations only; it is unlikely they could be deployed in real-world settings.

**Figure 5.**
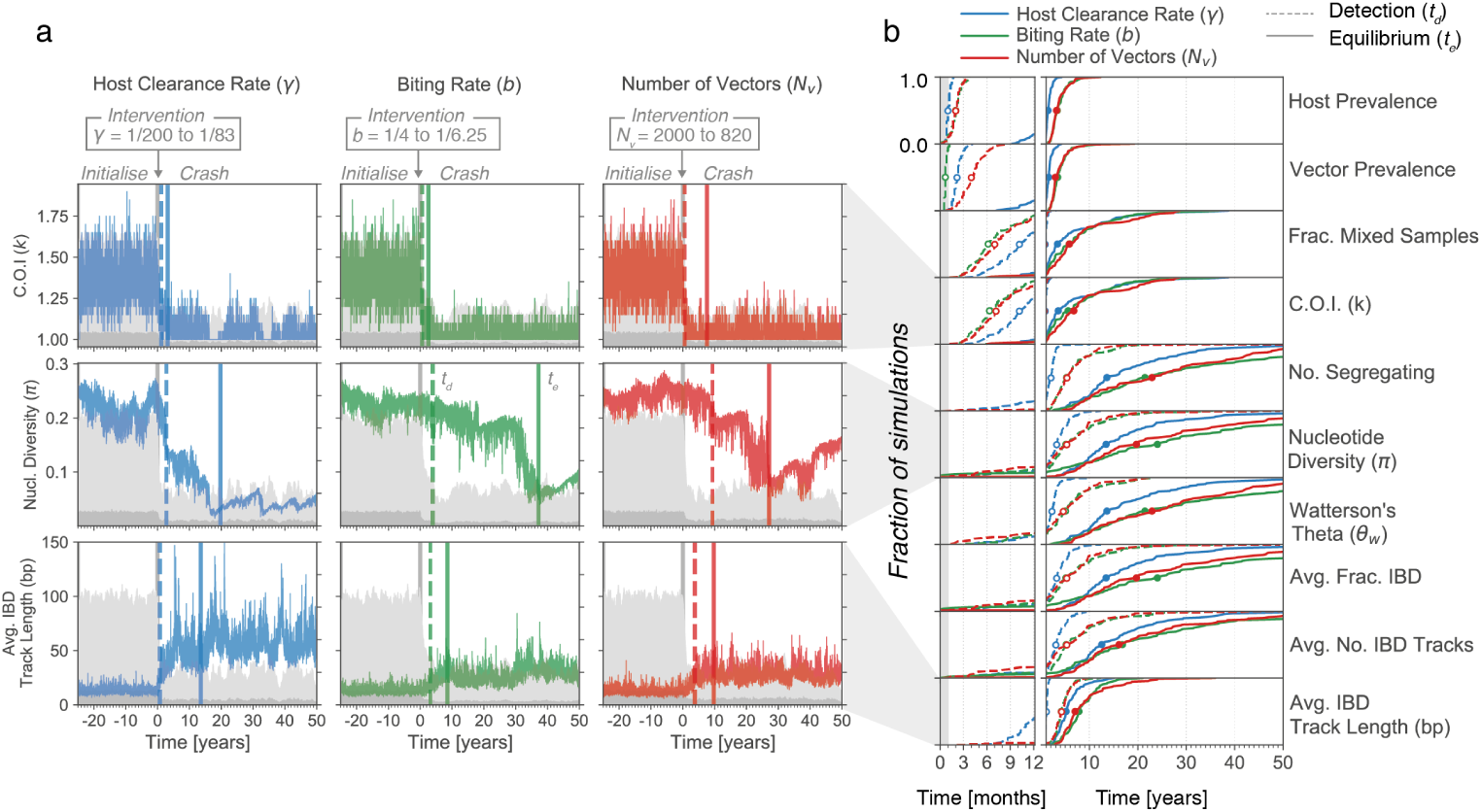
Detection and equilibrium times of genetic diversity statistics following a crash in parasite prevalence. (a) Each plot shows the behaviour of a genetic diversity statistic in an individual simulation through a crash in parasite prevalence, induced by: left column, increasing the host clearance rate (*γ*); middle column, reducing the vector biting rate (*b*); or right column, reducing the number of vectors. The intervention occurs at time zero (x-axis, grey vertical bar) in all cases. For each plot, the detection time (vertical dashed bar) and equilibrium time (vertical solid bar) of the genetic diversity statistic is indicated. Note that here a single simulation is shown for each intervention type. (b) Empirical cumulative density functions (ECDFs) of the detection and equilibrium times of diversity statistics, created from 100 independent replicate simulations for each intervention type. The y-axis gives the fraction of replicate simulations with a detection (dashed line) or equilibrium (solid line) less than the time indicated on the x-axis. Line color specifies the type of intervention. Open and closed circles give medians for the detection and equilibrium times, respectively. The first year is magnified for clarity.

Fig 5b shows the empirical cumulative density functions for *t*_*d*_ and *t*_*e*_ across all considered genetic diversity statistics, following the host prevalence decline at the beginning of the Crash epoch. Consistent with our observations from the averaged trajectories, the fastest responding statistics are those related to mixed infections: both the fraction of mixed samples and mean C.O.I. have a median detection time of between 6-10 months (depending on the epidemiological parameter causing the prevalence change). In the case of a *P f P R* decline resulting from an increase in host clearance rate, the average IBD track length had a median detection time of just over a year. All other detection times were in the range of 3-5 years. Similarly, the equilibrium times of mixed infection related statistics were fastest (median ≃ 3 − 6 years), followed by the average IBD track length (median ≃ 5 − 8 years). Strikingly, the median equilibrium times for many of the other statistics exceeded 20 years.

In terms of the relative behaviour of the statistics, the *t*_*d*_ and *t*_*e*_ values for the Recovery epoch were similar (S16 Fig), with mean C.O.I. and the fraction of mixed samples being the fastest statistics, and nucleotide diversity being the slowest. The median times tended to be longer, in particular for *t*_*e*_. This is likely a result of the rate of diversity being re-established (by mutation) being slower than the rate of it being eliminated (by an intervention).

#### Seasonality

We next aimed to explore whether any measures of genetic diversity were responsive to changes in parasite prevalence driven by seasonality. To this end, we developed a simulation framework where the number of vectors oscillates between a peak reached in the wet season and trough reached in the dry season, with *P f P R* fluctuating between ~0.6 and ~0.2 (see Materials and Methods). Vectors that die entering the dry season are selected at random, with the dry season lasting 170 days and the wet season lasting 195 days. Consistent with our results from the intervention analysis, we found that the fraction of mixed samples and mean C.O.I showed clear correlations with seasonal change in prevalence (*r*^2^ = 0.49 for mean C.O.I, *r*^2^ = 0.5 for fraction mixed samples; Fig 6 and S17 Fig). We found that the average IBD track length also exhibited a weak correlation with seasonally varying prevalence (*r*^2^ = 0.09). However, compared to equilibrium patterns, the relationship is in the opposite direction – with an increase in average IBD track length at higher prevalence values. This is likely driven by “epidemic expansion” in the early wet season – with the parasite population expanding faster than it acquires new mutations, resulting in increased IBD [30]. Consistent with this, we observed similar increases in average IBD track length in the first year of the Recovery epoch explored in the previous section (S20 Fig).

**Figure 6.**
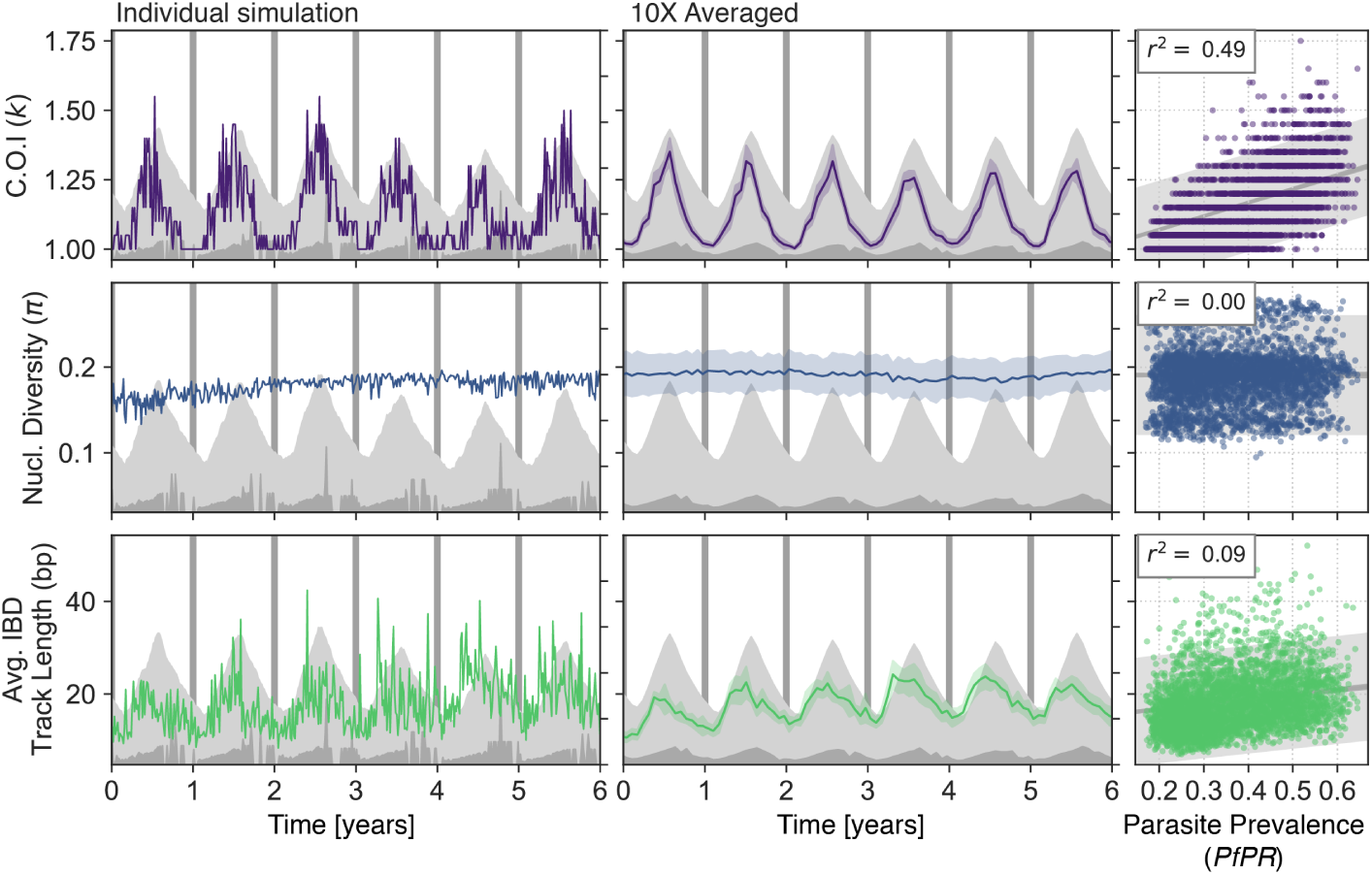
Responses of genetic diversity statistics to seasonal change in parasite prevalence. Annual variation in parasite prevalence was induced by varying the number of vectors (see Materials and Methods). The behaviour of genetic diversity statistics for an individual simulation is shown at left. The mean behaviour of 10 independent replicate simulations is shown at middle, with shaded areas giving the 95% confidence intervals. Scatterplots at right show the relationship between each genetic diversity at parasite prevalence across the six years of seasonal fluctuation. Each point represents a genetic diversity estimate (y-axis) computed from sampling parasite genomes from 20 infected hosts in an individual simulation; parasite prevalence (x-axis) is computed across the entire host population at the same time. Data from all 10 replicate simulations has been aggregated. Variance explained *r*^2^ from an ordinary linear regression is indicated at top left.

Notably, none of the other genetic diversity statistics we calculated exhibited correlations with seasonal fluctuation in prevalence (S17 Fig to S19 Fig).

## Discussion

The collection of parasite genetic data may, over the next few years, become a routine part of malaria surveillance. Yet, deriving the maximum benefit from this data will require an understanding of the relationships between malaria genetic diversity and epidemiology. Such understanding can be guided by modelling approaches, but only if both evolutionary and epidemiological processes are integrated. At present this is rare, as classical models in population genetics are poor approximations of the malaria life cycle, and classical epidemiological models don’t incorporate the evolution of parasites. To address this issue, we have combined the Ross-Macdonald and Moran models into a single framework, which we have implemented as a stochastic simulation called forward-dream.

We have used forward-dream to investigate the relationships between parasite genetic diversity and parasite prevalence in equilibrium and non-equilibrium settings. We find that many measures of parasite genetic diversity correlate with parasite prevalence at equilibrium. Our findings align with existing empirical data [25, 30-33], and support the idea that the rate of mixed infection (and as a consequence the rate of recombination) is positively correlated with parasite prevalence [34]. Moreover, we find that, for a given human host population size, statistics that reflect the long-term effective population size (*N*_*e*_) of the parasite, such as a nucleotide diversity and the number of segregating sites, also increase with equilibrium prevalence. We also explored the behaviour of these genetic diversity statistics in non-equilibrium settings, most importantly in response to changes in parasite prevalence that mimic malaria control interventions. Other authors have emphasised that the viability of a genomic approach to malaria surveillance will depend on how rapidly signals of epidemiological change become detectable in a reasonably sized sample of parasite genetic data [25]. We find that statistics related to the C.O.I distribution respond most rapidly (on the order of months), whereas other statistics, such as nucleotide diversity, may take decades to respond to a change in parasite prevalence.

Our results also demonstrate how relationships between prevalence and genetic variation are sensitive to the underlying epidemiological process. Specifically, we find that changes to the host clearance rate had a more profound effect on several genetic diversity statistics than changes to either the vector biting rate or density. The statistics exhibiting this behaviour are all related to *θ* = *N*_*e*_*µ*. As we did not alter the mutation rate across simulations, we expect this observation is being driven by effects on Ne . Where a population’s size fluctuates through time, the *N*_*e*_ can be approximated as the harmonic mean of those sizes, and thus is more influenced by periods where the population is small [35]. Similarly, a *P. falciparum* lineage alternates between a large vector population and a small host population, and so one explanation for the observation is that the amount of diversity is impacted more by changes influencing the smaller host population. This result complicates efforts to use genetic variation metrics to compare parasite prevalence across space or time, as it implies that only under certain conditions will changes in prevalence be reflected by changes in genetic variation.

forward-dream has several limitations, most obviously with respect to its simplicity. Many biological and epidemiological phenomenon are omitted, though they may have relevance to our results. For example, heterogeneous biting, acquired immunity, and migration are all phenomenon that have been proposed to influence the rate of mixed infections [2], yet they are currently not included in forward-dream. At present we do not explore the effect of selection, though this may be relevant in many contexts, particular in Southeast Asia where drug resistance is widespread [36]. Furthermore the genetic material simulated by forward-dream is equivalent to only a single chromosome harbouring one-thousand SNPs and, being neither infinite-sites nor infinite-alleles, we allow for reversible mutation, which influences some IBD statistics. Finally, the populations simulated in this study were small, typically with several hundred hosts and several thousand vectors. As a consequence, we were unable to explore parasite prevalence values below ~0.2, as the parasite population tended to go extinct before simulations reached equilibrium. Yet, as prevalence declines such regions are of particular interest, especially given the difficulties they produce with respect to estimating transmission intensity [12].

Many of the above limitations can be resolved with continued development of forward-dream. However, there are at least two salient considerations. The first is that additional complexity will likely increase computational costs. Merging the Moran and Ross-Macdonald models resulted in forward-dream being more computationally expensive than either, and for most of the simulations in this study run-times were 5-15 hours per experiment. Beyond optimizations to implementation and choice of programming language, there are some avenues by which efficiency could be improved. Reverse-time simulations harnessing coalescent theory can have greatly accelerated computational times, by omitting processes extraneous to the sample of genetic data collected (for example [37]), though the reverse-time formulation of the model described here is yet to be elucidated. In a similar vein, the development of models separating epidemiological and genetic processes are underway (see [38]), and could result in significantly faster simulations. Secondly, it is important to consider that more complex models typically require more parameters. Most likely these models will be both analytically intractable and statistically non-identifiable, thus making inference about their values impossible without additional and complex field experiments. Indeed, many of the parameters used within the current model have substantial uncertainty and were hard to find in current literature (see S1 Appendix). Community efforts to collate existing knowledge and address key uncertainties through experimental work would greatly benefit the field.

The value of our approach, as demonstrated here, is to use forward-dream as a tool through which relationships between genetics and epidemiology can be explored and experimental and analytical strategies can be evaluated. As methods for the inference of transmission intensity are developed, forward-dream can provide a basis for assessing their expected performance, and designing ideal sampling strategies under different epidemiological scenarios. It is in these ways that forward-dream, and future simulations like it, can provide a platform for interpreting the signals within the projected tens of thousands of malaria genomes that will be collected over the next decade, and can help to leverage those signals for malaria surveillance.

## Materials and methods

### Availability

All of the code developed as part of this manuscript, including forward-dream, is available on GitHub (https://github.com/JasonAHendry/fwd-dream).

### Collection of genetic data and computation of summary statistics

For all of the simulations in this manuscript, parasite genomes were collected from randomly selected infected hosts. For each host, we simulated DNA sequencing by taking a subset of all parasite genomes within the host 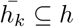 (where 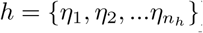) such that each genome in 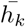 was different from all others at a minimum of 5% of its sites; the assumption being that genomes more similar than this would not be readily distinguishable by sequencing. The C.O.I. of each host is then 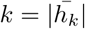.

The fraction of mixed samples and the mean C.O.I. is computed directly from the distribution of *k* across all sequenced hosts. To compute other statistics, we pooled all genomes collected across all hosts. The number of segregating sites, the number of singletons, nucleotide diversity, Watterson’s Theta (*θ*_*w*_), and Tajima’s *D* were calculated using scikit-alllel (https://scikit-allel.readthedocs.io/en/stable/). We estimated identity-by-descent (IBD) profiles between pairs of parasite genomes using identity-by-state (IBS). Since the genetic layer of forward-dream is not an infinite alleles model, and mutation is reversible, this leads to an inflation of IBD statistics; trends in IBD are preserved.

### Averaging of genetic diversity statistics across independent forward-dream simulations

To produce smoothed trajectories of genetic diversity statistics for the intervention analysis (Fig 4b, d and S12 Fig to S14 Fig), we averaged independent replicate forward-dream simulations. As forward-dream operates in continuous-time, parasite genetic data is never sampled at exactly the same time in independent simulations. Thus, to average simulations, we binned time into 25-day intervals. Finally, across all replicate simulations and for the entire duration of the intervention analysis, genetic diversity statistics computed within each 25-day bin were averaged.

### Computing response time statistics *t*_*d*_ and *t*_*e*_

We created two simple metrics to characterize the temporal response of different genetic diversity to changes in *P f P R*. These metrics were developed specifically in the context of the intervention experiments described in the Results section, and they are computed for individual simulations. The “detection time” (*t*_*d*_) estimates the amount of time before a change in parasite prevalence would be detected, if that change was being monitored for using a given genetic diversity statistic. To compute it, we first construct a distribution for the genetic diversity statistic of interest at equilibrium. When considering a decline in parasite prevalence, this is achieved by recording the genetic diversity statistic’s value for 25 years proceeding the Crash epoch, during which time the simulation is at equilibrium (at a host prevalence value of 0.65). For an increase, the statistic is recorded for 25 years proceeding the Recovery epoch. *t*_*d*_ is then computed as the first time, after the prevalence chance has occurred, that three consecutive samples have a value for that statistic outside of the quantile interval [*α*/2, 1 − *α*/2], with *α* = 0.01; i.e. the first time when three consecutive samples have a value that would be observed with a probability of less than 1% if the simulation were at equilibrium. Requiring that three consecutive samples (equivalent to approximately two weeks of genetic data) have values outside the interval makes *t*_*d*_ more robust to the high variability observed in individual simulations.

Similarly, the “equilibrium time” (*t*_*e*_) estimates the time until a given genetic diversity statistic regains equilibrium following a host prevalence change. *t*_*e*_ is computed as the first time that six consecutive samples have a value within the inter-quartile range ([*α*/2, 1 − *α*/2], with *α* = 0.5), of the distribution of the statistic at its new equilibrium. Again, requiring six consecutive values within the inter-quartile range makes our estimates of *t*_*e*_ more robust to the high variability of individual simulations; we elected for six rather than only three samples as the criterion of being within the inter-quartile range is weaker than the *t*_*d*_ criterion.

### Parameters for malaria control intervention and seasonality experiments

The complete parameter files (stored as ‘.ini’) used to specify the malaria control intervention and seasonality experiments are available on GitHub within the ‘params’ directory. All of these experiments began with the same set of parameters listed in Table 1 before individual parameters were changed to either mimic malaria control interventions or induce seasonality. To achieve a parasite prevalence of 0.2 during the Crash epoch of malaria control intervention experiments, either the host clearance rate (*γ*) was increased to 0.012, the vector biting rate (*b*) was reduced to 0.16, or the number of vectors (*N*_*v*_) was reduced to 819. In the Recovery epoch they were returned to their original values. To achieve an annually varying parasite prevalence in the seasonality experiment, the number of vectors (*N*_*v*_) was oscillated between 10 during a dry season lasting 170 days, and 2800 in the wet season lasting 195 days.

## Supporting information

**S1 Appendix. Additional information on forward-dream implementation and parameterisation.**

**S1 Fig. Validating equilibrium host prevalence values in forward-dream.** The epidemiological layer of forward-dream implements the Ross-Macondald model, where the host prevalence is a function of the rate parameters (see Eq. 1). Violinplots summarize the prevalence values observed in forward-dream simulations with expected equilibrium prevalence values varying from 0.2 to 0.8 (computed using Eq. 1) given on the x-axis. The different equilibrium prevalence values were achieved by varying either the host clearance rate (*γ*), the vector biting rate (*b*), or the number of vectors (*N*_*v*_). The variance explained (*r*^2^) in an ordinary linear regression is shown at top-left of each plot.

**S2 Fig. Validating intra-host fixation times in forward-dream.** (a) The infection of a single host is evolved through time and the within-host alelle frequency of a given site is indicated by the red line. The site fixes around day 125. The experiment is repeated 1000 times (grey lines) and the fraction of infections fixed at a given time is indicated by the blue line. All experiments started with an initial allele frequency of 0.5 and a drift rate of 1/event per day. (b) Distribution of fixation times from (a). The observed mean (64.81 days) is very close to the theoretically expected mean from the Moran model (64.56 days). (c) The experiment in (a) is repeated but with different initial allele frequencies (x-axis) and three different drift rates (light blue, dark blue, and green line). In all cases, the observed mean fixation times are close to the theoretically expected times. Shading gives 95% confidence intervals for mean estimates.

**S3 Fig. Varying equilibrium prevalence values in forward-dream.** The parameter values of forward-dream are varied to produce simulations with equilibrium parasite prevalence values varying from 0.2 to 0.8. (a) Varying the number of vectors. Prevalence in hosts indicated in red, vectors in blue. Dots mark parasite prevalence values of 0.1 through 0.8. (b) Varying the vector biting rate *b*. Note 1*/b* gives the average time between successive bites, show in right plot. (c) Varying the host clearance rate (*γ*). Note 1/*γ* gives the average duration of host infection, shown in right plot.

**S4 Fig. Equilibrium relationships between parasite prevalence and mixed infection related statistics.** Distributions of mixed infection related genetic diversity statistics (y-axis), plotted for equilibrium parasite prevalence values tuned to between 0.2 and 0.8 (x-axis) in forward-dream simulations. Left, middle, and right columns show distributions when parasite prevalence is varied as a function of the host clearance rate (*γ*, in blue), vector biting rate (*b*, in green) or number of vectors (*N*_*v*_, in green). Each boxplot contains the result of 30 replicate experiments, where the parasite genomes within 20 randomly selected hosts are collected at every 30 days for 10 years and are used to compute the genetic statistic of interest. The variance explained by ordinary least squares regression is given at top left, and line of best fit and confidence intervals indicated in grey.

**S5 Fig. Equilibrium relationships between parasite prevalence and genetic diversity statistics related to the size and shape of the sample genealogy.** See S4 Fig for details.

**S6 Fig. Equilibrium relationships between parasite prevalence and genetic diversity statistics related to identity-by-descent patterns.** See S4 Fig for details.

**S7 Fig. Co-linearity between different genetic diversity statistics in forward-dream.** Matrices of Pearson’s Correlation Co-efficient (*R*) calculated between all pairs of genetic diversity statistics is shown. In panel (a) host prevalence was tuned to different values between 0.2 and 0.8 by varying the host clearance rate (*γ*); (b) by varying the vector biting rate (*b*); or (c) by varying the number of vectors *N*_*v*_. In all cases there is significant co-linearity between different genetic diversity statistics.

**S8 Fig. Behaviour of an individual forward-dream simulation during a crash and recovery of parasite prevalence: driven by a change in host clearance rate (*γ*).** (a) Prevalence (y-axis) over a 220-year period where a population crash and recovery has occurred. The simulation is first allowed to equilibrate (”Initialise” Epoch). At time zero (x-axis), the host clearance rate is increased from 0.005 (1/200) to 0.012 (1/83), causing a crash in prevalence (”Crash” Epoch). After 125 years (sufficient time to for the simulation to reach a new equilibrium), the host clearance rate is returned to its original value (”Recovery” Epoch). Prevalence of all infected (*k* > 0) and multiply-infected (*k* > 1) hosts is indicated by red and pink lines, respectively. The same is shown for vectors in blues. (b) The same simulatoin as in (a), but with a variety of genetic diversity statistics shown. Note that the statistics are computed from parasite genomes collected from 20 randomly selected hosts every 5 days. For reference, light and dark grey show the host and vector prevalence on the second y-axis.

**S9 Fig. Behaviour of an individual forward-dream simulation during a crash and recovery of parasite prevalence: driven by a change in vector biting rate (*b*).** See S8 Fig for details.

**S10 Fig. Behaviour of an individual forward-dream simulation during a crash and recovery of parasite prevalence: driven by a change in number of vectors (*N*_*v*_).** See S8 Fig for details.

**S11 Fig. Temporal fluctuations in genetic diversity statistics of different frequencies, without any change in parasite prevalence.** Panel (a) shows the noisy behaviour of three genetic diversity statistics (y-axis) from an individual simulation where parasite prevalence was kept fixed at 0.65 for a 25 year period (x-axis). The trajectory of each statistic was smoothed using a rolling mean, with window sizes varying from 1 day (1 d., purple), which is equivalent to no smoothing, up to 10 years (10 y., yellow). The mean of the statistic during the 25-year window is indicated with the grey horizontal bar. Notice how even with a 10-year window rolling mean, the nucleotide divesity still deviates from its mean value. (b) Across 100 independent replicate simulations, the reduction in variance of each genetic diversity statistic with increasing rolling mean window sizes is shown. The y-axis gives the ratio of the variance for the window size indicated by the x-axis (*V ar*(*X*_*w*_)) divided by the unsmoothed variance in the genetic diversity statistic (*V ar*(*X*)). Increasing with window size of the rolling mean always reduces the variance, but at different rates for different statistics.

**S12 Fig. Average behaviour of 100 replicate forward-dream simulations during a crash and recovery of parasite prevalence: driven by a change in host clearance rate (*γ*).** Same as Fig S8 Fig, but instead of showing an individual simulation trajectory, the averaged trajectory of 100 replicate simulations is shown. Each line is a mean across the 100 replicates, and the shading gives the 95% confidence intervals.

**S13 Fig. Average behaviour of 100 replicate forward-dream simulations during a crash and recovery of parasite prevalence: driven by a change in vector biting rate (*b*).** Same as Fig S9 Fig, but instead of showing an individual simulation trajectory, the averaged trajectory of 100 replicate simulations is shown. Each line is a mean across the 100 replicates, and the shading gives the 95% confidence intervals.

**S14 Fig. Average behaviour of 100 replicate forward-dream simulations during a crash and recovery of parasite prevalence: driven by a change in the number of vectors (*N*_*v*_).** Same as Fig S10 Fig, but instead of showing an individual simulation trajectory, the averaged trajectory of 100 replicate simulations is shown. Each line is a mean across the 100 replicates, and the shading gives the 95% confidence intervals.

**S15 Fig. Average behaviour of Tajima’s *D* during a crash and recovery in parasite prevalence.** Same data as Figures S12 Fig-S14 Fig. Colored lines show mean estimate across 100 replicate simulations, shaded area gives 95% confidence intervals. Notice how Tajima’s *D* increases during a population contraction and decreases during population growth.

**S16 Fig. Detection and equilibrium times of genetic diversity statistics following a recovery in parasite prevalence.** (a) Each plot shows the behaviour of a genetic diversity statistic in an individual simulation through a recovery of parasite prevalence, induced by: left column, reducing the host clearance rate (*γ*); middle column, increasing the vector biting rate (*b*); or right column, increasing the number of vectors. The intervention occurs at time zero (x-axis, grey vertical bar) in all cases. For each plot, the detection time (vertical dashed bar) and equilibrium time (vertical solid bar) of the genetic diversity statistic is indicated. Note that here a single simulation is shown for each intervention type. (b) Empirical cumulative density functions (ECDFs) of the detection and equilibrium times of diversity statistics, created from 100 independent replicate simulations for each intervention type. The y-axis gives the fraction of replicate simulations with a detection (dashed line) or equilibrium (solid line) less than the time indicated on the x-axis. Line color specifies the type of intervention. Open and closed circles give medians for the detection and equilibrium times, respectively. The first year is magnified for clarity.

**S17 Fig. Responses of mixed infection related genetic diversity statistics to seasonal change in parasite prevalence.** Annual variation in parasite prevalence was induced by varying the number of vectors. The behaviour of genetic diversity statistics for an individual simulation is shown at left. The mean behaviour of 10 independent replicate simulations is shown at middle, with shaded areas giving the 95% confidence intervals. Scatterplots at right show the relationship between each genetic diversity at parasite prevalence across the six years of seasonal fluctuation. Each point represents a genetic diversity estimate (y-axis) computed from sampling parasite genomes from 20 infected hosts in an individual simulation; parasite prevalence (x-axis) is computed across the entire host population at the same time. Data from all 10 replicate simulations has been aggregated. Variance explained *r*^2^ from an ordinary linear regression is indicated at top left.

**S18 Fig. Responses of sample genealogy related genetic diversity statistics to seasonal change in parasite prevalence. See S17 Fig for details.**

**S19 Fig. Responses of IBD related genetic diversity statistics to seasonal change in parasite prevalence.** See S19 Fig for details.

**S20 Fig. Zoom on response of average IBD track length to increase in prevalence at beginning of Recovery epoch.** Same data as S12 Fig to S14 Fig but focussing only on the average IBD track length and a four-year window around the beginning of the recovery. Notice how for a change in the number of vectors (*N*_*v*_) there is an increase in average IBD track length at the beginning of the recovery, consistent with epidemic expansion.

**S1 Table. Genetic diversity statistics computed in forward-dream.**

## Acknowledgments

This study was supported by the Wellcome Trust (206194, 090770, 204911, 100956/Z/13/Z to GM, and 109107/Z/15/Z to JH) and the Li Ka Shing Foundation (to GM). The computational aspects of this research were supported by the Wellcome Trust Core Award Grant Number 203141/Z/16/Z and the NIHR Oxford BRC. The views expressed are those of the author(s) and not necessarily those of the NHS, the NIHR or the Department of Health. Many thanks to Tim Anderson and Lisa J. White for helpful discussions.

