## Supplementary Information for "Elucidating relationships between *P.falciparum* prevalence and measures of genetic diversity with a combined genetic-epidemiological model of malaria"

### Additional details on the model structure of `forward-dream`

Jason A. Hendry

May 2020

#### 1 Implementation

In `forward-dream`, time is continuous and the occurrence of events is dictated by a competing Poisson process. The rate at which any event occurs, $r_{total}$, is equal to:

$$r_{total} = bN_v + \gamma h_1 + \epsilon v_1$$

which is the sum of the total biting rate across all vectors, the clearance rate across all infected hosts and the clearance rate across all infected vectors. The time until the next event is drawn from an exponential distribution with an expectation of $1/r_{total}$. Conditional on an event occurring, the type of event is sampled proportional to its fractional rate. For example, the probability of a given event being a biting event is:

$$P(bite|event) = \frac{bN_v}{r_{total}}$$

If a biting event occurs, the host and vector involved are sampled at random from the population, and the nature of parasite transmission depends on their infection status. Similarly, for clearance events, the host or vector to be cleared of infection (i.e. returned to the susceptible state) is chosen at random.

#### 2 Parameterisation

`forward-dream` is completely specified by 17 parameter values (see Table 1 of the main text). Below we discuss how these parameters were set for the simulations presented in this study.

##### Biting rate ($b$)

The biting rate ($b$) gives the daily rate at which a single vector bites a host, such that $1/b$ gives the average time between bites. It is typically stated that $1/b$ ranges from 2-4 days, and common in modelling papers is to give an assumed value of $1/b = 3$. This is supported by some laboratory studies; for example [19] observed an approximately 3 day feeding and oviposition behaviour of *Anopheles gambiae* when offered daily blood meals and kept alone.

In `forward-dream` we set the default biting rate $b = 0.25$ which implies an average of 4 days between bites. We elected for a slightly lower default biting rate to partially compensate for the absence of an incubation time within the vector; bringing the average number of infectious bites delivered by a vector closer to what it would be were an incubation time included.

##### Vector-to-host ($\pi_h$) and host-to-vector ($\pi_v$) transmission efficiencies

The vector-to-host transmission efficiency ($\pi_h$) is the probability that a host becomes infected after being bitten by an infectious vector. An assembly and reanalysis of multiple sources of information on $\pi_h$ has been conducted by [34]. The most direct source of information on transmission efficiency they identify comes from the control arm of drug or vaccine trials, where immunologically naive individuals are challenged with mosquitoes carrying *P. falciparum* [34]. Using data of this kind, [34] estimate a vector-to-host transmission efficiency of 55%. A second source of information comes from simultaneously collecting data on the force of infection ($FOI$, the rate at which hosts become infected per unit time) and the entomological inoculation rate ($EIR$, the average number of infectious bites received per host per unit time). The ratio $\frac{FOI}{EIR}$ is then taken to equal the vector-to-host transmission efficiency [34]. A study of this type conducted in Saradidi, Kenya, estimated the vector-to-host transmission efficiency in children to be 7.5% [6]. Reanalysis by [34], fitting three different mathematical models (unrooted linear, linear, and a model accommodating for heterogeneous biting) to observations of $\frac{FOI}{EIR}$ collected through time, produced estimates of 5%, 2% and 27% (for the three models, respectively). A key conclusion of [34] was that transmission efficiency declines at higher transmission intensities (in particular when $EIR > 10$), and that this trend can be fit well using a heterogeneous biting model with $\pi_h = 0.55$. However, when models without heterogeneous biting are used, $\pi_h = 0.55$ produces $FOI$ values that are significantly above observations; lower values of transmission efficiency provide a better fit to the data.

As `forward-dream` does not model heterogeneous biting, we set the default vector-to-host transmission efficiency to 10% ($\pi_h = 0.1$), roughly in the middle of the raw estimates reported by [34].

The host-to-vector transmission efficiency ($\pi_v$) is the probability that a vector becomes infected after biting an infectious host. Analysing 23-years of longitudinal data collected on the prevalence of infection in both hosts and vectors in Dielmo, Senegal, [12] produced estimates of $\pi_v$. Similarly to the vector-to-host transmission efficiency, [12] found that the $\pi_v$ was lower in periods of higher transmission: ranging from $\sim 0.18$ at a parasite prevalence ($PR$) of 0.2, to $\sim 0.03$ at $PR = 0.7$ (for *Anopheles gambiae s.l.*).

In `forward-dream` we set the default host-to-vector transmission efficiency to 10% ($\pi_h = 0.1$).

##### Host infection clearance rate ($\gamma$)

The host infection clearance rate, $\gamma$, is the daily rate at which hosts in the infected state ($H_1$) return to the susceptible state ($H_0$). The host infection clearance rate is modelled as a constant, and so the duration of host infection is exponentially distributed with a mean of $1/\gamma$. Recall that in `forward-dream`, both singly- and multiply infected hosts are in the $H_1$ state. As $\gamma$ is the same for all hosts in the $H_1$ state, multiple infection has no effect on the duration of host infection in `forward-dream`; all hosts have the same daily rate of clearance $\gamma$, regardless of the composition of their infection.

| Estimate (days) | Study Type | Notes | Reference |
|---|---|---|---|
| 211.6 | Malariatherapy | | [30] |
| 54.9 | *msp2* analysis | Gamma fit | [9] |
| 139.9 | *msp2* analysis | Weibull fit | [9] |
| 205.3 | *msp2* analysis | lognormal fit | [9] |
| 219.7 | *msp2* analysis | Exponential fit | [9] |
| 152 | *msp2* analysis | | [31] |
| 155 | *msp2* analysis | <1 yo. | [18] |
| 256 | *msp2* analysis | 1-2 yo. | [18] |
| 257 | *msp2* analysis | 3-4 yo. | [18] |
| 319 | *msp2* analysis | 5-9 yo. | [18] |
| 176 | *msp2* analysis | 10-19 yo. | [18] |
| 129 | *msp2* analysis | 20-39 yo. | [18] |
| 126 | *msp2* analysis | 40-59 yo. | [18] |
| 131 | *msp2* analysis | >59 yo. | [18] |

Table 1: A selection of estimates of the duration of *P. falciparum* infection. Estimates from [9, 31, 18] are all from the study of *msp2* alleles in Northern Ghana, but analyzed using different methods. In [18] estimates were aged-stratified. Horizontal lines separate estimates from individual studies. Abbreviations: yo., years old.

Studies examining the duration of untreated *P. falciparum* infections are scarce on account of the medical obligation to treat [30]. In the mid-1900s in the USA, the recommended treatment for individuals with neurosyphilis was deliberate infection with *P. falciparum* malaria, and analysis of the progression of these infections has become an important source of information about untreated infection duration. Seeking to understand both the average duration of untreated infection and the best fitting distributional form, [30] analysed data from 54 such malariatherapy patients who were inoculated with *P. falciparum* either by sporozoites or infected blood, and were subsequently followed by daily microscopic examination. They observed a mean infection duration of 211.6 days [30]. It is also notable that they found that both the Gompetz and Weibull distributions provided better fits than the exponential model [30]. This data source has significant caveats, however, including that the adults involved were all naïve and co-infected with syphilisis, and the strains used were selected for having a low clinical virulence and the greatest curative properties [25]. Ultimately, the extent to which these infections reflect those developed in natural settings is unclear.

A second source of data on untreated infections comes from a study conducted in the Kassena-Nankana District of Northern Ghana in 2000-2001, a region with very high transmission ($EIR > 300$) [29]. In this study, approximately 300 asymptomatic individuals positive for malaria were tracked for a ten month period, where every two months blood samples were taken and both microscopic and PCR analysis of the *msp2* gene was performed [29]. The analysis of the *msp2* gene allowed for the tracking of individual parasite genotypes in a context where most individuals were multiply infected. A variety of authors have analysed this data source using different methods, and estimates of the duration of infection range from as low as 55 to as high as 319 days (see Table 1).

The review by [2] highlights other sources of information on infection duration, including accidental cases of malaria infection by blood transfusion, and provides evidence that in some cases a *P. falciparum* can last for multiple years.

For `forward-dream`, we set the default host infection clearance rate $\gamma = 0.005$, corresponding to an average duration of infection of 200 days. This is Macdonald's original estimate of infection duration [24], and broadly consistent with the estimates described above.

##### Vector infection clearance rate ($\epsilon$)

In `forward-dream`, the vector infection clearance rate, $\epsilon$, is the daily rate at which vectors in the infected state ($v_1$) return to the susceptible state ($v_0$). In the Ross-Macdonald model, it is classically assumed that vectors do not clear infections, but rather that the flow from $v_1$ to $v_0$ is driven by the death of infected vectors and their (instantaneous) replacement by susceptible vectors (reviewed in [33]). This allows for the vector population size ($N_v$) to be kept fixed. It further implies that $\epsilon$ represents the daily probability of death for infected vectors. As $\epsilon$ is a constant, the duration of vector lifespans is exponentially distributed with a mean of $1/\epsilon$. Consequently, vectors do not have an increased probability of death with age (they do not experience senescence); the common justification for this is that mosquitoes die too rapidly of other causes to die as a consequence of aging.

To set $\epsilon$, we looked for data on either lifespan or the daily survival rate (commonly denoted $p$) of *Anopheles* mosquitoes from the field. Since we are looking to parameterise infected mosquitoes, we are particularly interested in data on female *Anopheles* mosquitoes. We note that there are a large variety of factors that influence mosquito survival rate in nature – including the climate/weather, most pertinently the temperature and humidity, the species and activities of the mosquito, and the presence of parasites and predators – and that these factors likely vary substantially through time and across different geographies [3]. Thus, in malaria endemic areas there is likely variation in $\epsilon$ through space in time.

A relatively straightforward approach to estimating the lifespan of *Anopheles* mosquitoes is by breeding them in controlled laboratory environments or outdoor cages. However, estimates of this kind are generally thought to be inflated, as many common sources of mortality that would act on wild vectors are not in effect [11]. [19] found that the maximum lifespan of a laboratory colony *A. gambiae* mosquitoes, fed on a 10% sucrose solution, was 16 days at 27°C and 34 days at 22°C ($p = 0.94$ and $p = 0.97$, respectively). Given that mortality in laboratory experiments is driven primarily by senescence, whereas in the Ross-Macdonald model it is implicitly assumed that mortality due to senescence is negligible, we consider these estimates unsuitable for parameterising `forward-dream`.

In general, the lifespan or survivorship of wild vectors cannot be measured directly [3]. However, a multitude of methods have been developed to provide indirect estimates. Below we briefly outline two general types of estimation approaches, and in Table 2 we provide a selection of estimates arising from them.

For a more detailed discussion and additional estimates, the reader should consult Chapter 13 of [32] which covers the subject extensively.

| $p$ | Method | Location | Species | Details | Ref. |
|---|---|---|---|---|---|
| 0.74 | Mark-release-recapture | Burkina Faso | *An. gambiae* | | [16] |
| 0.77 | Age-composition | Tanzania | *An. gambiae* | Michenga | [10] |
| 0.78 | Age-composition | D.R.C. | *An. gambiae* | Urban | [14] |
| 0.81 | Age-composition | Senegal | *An. arabiensis* | | [35] |
| 0.82 | Age-composition | Kenya | *An. gambiae* | | [11] |
| 0.83 | Age-composition | Nigeria | *An. gambiae* | | [1] |
| 0.84 | Age-composition | Tanzania | *An. gambiae* | Namawala | [10] |
| 0.84 | Mark-release-recapture | Tanzania | *An. gambiae* | | [20] |
| 0.87 | Age-composition | Kenya | *An. funestus* | | [11] |
| 0.91 | Age-composition | Tanzania | *An. gambiae* | | [17] |
| 0.91 | Age-composition | D.R.C. | *An. gambiae* | Rural | [14] |
| 0.93 | Age-composition | Uganda | *An. gambiae* | | [17] |

Table 2: A selection of daily survival rate ($p$) estimates for *Anopheles* vectors. Note that in `forward-dream` the vector infection clearance rate is $\epsilon = 1 - p$.

The first type of approach is based on Mark-release-recapture experiments. Here, mosquitoes are marked, originally with paint or a radioisotope, and released in a natural environment [20]. It has been observed that the number of marked mosquitoes recaptured on every subsequent day declines, and it assumed this is due to mortality [20]. From this decline, daily mortality can be estimated in a variety of ways (reviewed in [32]). This type of approach can be challenged by low recapture numbers, changes in mosquito survival or behaviour on account of the marking, and the confounding of mortality with other reasons for mosquito loss, such as dispersal.

A second approach looks for indicators of the age-composition of the mosquito population, typically with respect to developmental stages. For example, whether a mosquito is parous (has laid eggs) or nulliparous (has not laid eggs) can be estimated by dissection [17]. If an assumption is made about the duration of time until egg laying, the daily survival rate can be estimated by the proportion parous [17]. A great diversity of methods of this type exists, focusing different age-composition indicators (reviewed in [32]). In general, they all involve making assumptions about (and are sensitive to) the duration of the developmental stage(s) under consideration.

Overall, estimates of the survival rate ($p$) of *Anopheles* mosquitoes typically fall between 0.75 and 0.95 (see Table 2). For `forward-dream`, the vector infection clearance rate is $\epsilon = 1 - p$ and we set the default value at 0.2, corresponding to a survival rate of 0.8.

##### Number of sub-compartments in hosts ($n_h$) and vectors ($n_v$)

An individual host can harbour up to $10^9$-$10^{11}$ *P. falciparum* parasites [36]. However, in all the simulations described here, $n_h$ (and $n_v$) are set to ten. This is motivated primarily by reducing computational costs, but the simplification can also be justified biologically. First, even if an individual host contains $10^9$-$10^{11}$ parasites, *P. falciparum* replicates clonally within the host and so the majority of these will have near-identical genomes. A host may, however, be infected by multiple distinct parasite strains, generating a mixed infection. Importantly, setting $n_h = 10$ still accommodates for this possibility. Second, parasite genomes that exist at a frequency much less than 1/10 of the population are unlikely to produce substantial signal in genetic data. For example, with whole-genome sequencing data, sequencing depths are typically $\leq$ 100X and so parasites comprising less than 1/10 of the infection will be represented by only a handful of reads.

##### Mutation and drift rates in the host ($\theta_h$, $d_h$) and vector ($\theta_v$, $d_v$)

We have combined the mutation and drift rate parameterisation into a single section because in `forward-dream` mutational events occur conditional on drift events, and thus the parameters are related. In particular, considering hosts, drift events occur at a fixed constant rate $d_h$ such that for any time interval $\delta t$ the number of drift events will be drawn from $\sim Poi(\delta t d_h)$. Note that `forward-dream` implements a Moran model, such that for each drift event two sub-compartments are selected at random, and one is duplicated while the other is deleted. With probability $\theta_h N_{snps}$, each of these events will be instead a mutational event. The same occurs for vectors with rates $d_v$ and $\theta_v$.

The malaria parasite genome in `forward-dream` is designed to represent a collection of SNPs, and there are at least two sources of information on the single-nucleotide variant (SNV) mutation rate of *P. falciparum*. The first is from clone-tree experiments, where *P. falciparum* strains are cultured *in vitro* and accumulated mutations are detected by whole-genome sequencing. From these experiments several estimates of the mutation rate of *P. falciparum* during the erythrocytic phase have been produced, typically on the order of $\sim 10^{-10}$ [13, 21]. A second approach is to look at sequence divergence from a related *Plasmodium* species, such as *Plasmodium riechenowi* [23]. Here, the estimated mutation rate corresponds to the entire generation of *P. falciparum*, i.e. including both the host and vector phases. We have tabulated a selection of these mutation rates below. We are unaware of any mutation rate estimates specific to the vector phase.

| SNV Rate | Value | Strain | Method | Ref. |
|---|---|---|---|---|
| per gen. per bp | 6.88 x $10^{-10}$ | Wild | Phylogenetic cf. to *P. riechenowi* | [23] |
| per gen. per bp | 4.90 x $10^{-10}$ | Wild | Phylogenetic cf. to *P. riechenowi* | [23] |
| per ieg per bp | 4.07 x $10^{-10}$ | 3D7 | Clone-tree | [13] |
| per ieg per bp | 3.63 x $10^{-10}$ | Dd2 | Clone-tree | [13] |
| per ieg per bp | 3.78 x $10^{-10}$ | HB3 | Clone-tree | [13] |
| per ieg per bp | 2.10 x $10^{-10}$ | 3D7 | Clone-tree | [21] |
| per ieg per bp | 3.20 x $10^{-10}$ | Dd2 | Clone-tree | [21] |
| per ieg per bp | 2.95 x $10^{-10}$ | HB3 | Clone-tree | [21] |
| per ieg per bp | 2.27 x $10^{-10}$ | KH-01 | Clone-tree | [21] |
| per ieg per bp | 1.64 x $10^{-10}$ | KH-02 | Clone-tree | [21] |
| per ieg per bp | 1.70 x $10^{-9}$ | 3D7 | Clone-tree | [8] |
| per ieg per bp | 3.20 x $10^{-9}$ | Dd2 | Clone-tree | [8] |

Table 3: A selection of mutation rate estimates for *P. falciparum*. The two different estimates from [23] arise from differing assumptions about the speciation time between *P. falciparum* and *P. reichenowi* (5Mya or 7Mya.). Note that [8] estimates are an order of magnitude higher, but have been adjusted to include unobserved lethal mutations.

It was unclear to us how to use these mutation rates to parameterise `forward-dream`. Given that $n_h = 10$ and $n_v = 10$, each sub-compartment can be thought of as representing the consensus sequence of 1/10th of the parasite population, rather than a single genome. This implies that a mutational event in `forward-dream` is equivalent to an event whereby a novel variant reaches a frequency 1/10, within the host/vector. Note that this is not co-incident with the mutation occurring, but is related also to the rate of drift. In addition, though $n_h$ and $n_v$ are fixed in `forward-dream`, in reality the size of the parasite population fluctuates dramatically in both the vectors and the hosts over the course of an infection – going through exponential growth phases and bottlenecks in response to immune pressure.

For simplicity, we set the host and vector mutation and drift rates equally, to $d_h = 1$, $d_v = 1$ and $\theta_h = 0.0001$, $\theta_v = 0.0001$. Thus, for both hosts and vectors, a drift event occurs on average every day and, with $N_{snps} = 1000$, one in every ten drift events produces a mutation. Since there are ten sub-compartments in both the hosts and the vectors, this implies that an individual sub-compartment acquires a mutation once every one-hundred days. In the future, estimates of these parameters can be strengthened by the collection of longitudinal WGS data for *P. falciparum*.

**Probability a given parasite genome passes through the host-to-vector ($p_h$) or vector-to-host bottleneck ($p_v$)**

The two parameters $p_h$ and $p_v$ control the size of the bottleneck when parasites are being transmitted from the host to the vector, or vector to the host, respectively. In terms of the `forward-dream` model, $p^v$ gives the probability that a given sub-compartment in the vector is selected to colonize a host sub-compartment (see main text). Biologically, we expect it to be determined by the processes that reduce the amount of genetic diversity found in the initial blood-stage infection, relative to that present in the salivary glands of the vector that initiated that infection. This would include the number of sporozoites that are transmitted to the host, how representative they are of the genetic diversity within the salivary glands, and what fraction of them succeed in invading hepatocytes.

By dissecting wild caught *Anopheles gambiae* salivary glands and quantifying the abundance of sporozoite-specific antigens (either with an ELISA or immunoradiometric assay), the average number of sporozoites within an infectious vector has been found to be around 4000 to 5000, although the distributions are highly variable with some mosquitoes found carrying tens of thousands of sporozoites [5, 15]. Estimates of the number of sporozoites delivered in a single bite are also variable. Using a membrane feeding assay, [4] found that a geometric mean of 4.9 sporozoites were delivered by *An. freeborni* and 11.3 sporozoites were delivered by *An. gambiae* mosquitoes. In rodent model systems, two studies found an average of 123 and 281 *Plasmodium berghii* sporozoites were injected in a single bite, though again the distributions were highly variable with over a thousand sporozoites delivered in some cases [26, 22]. Cumulatively these results suggest that, typically, only a small percentage of salivary gland sporozoites are delivered to the host [4]. Finally, analysing early blood stage infections, [7] deduced that roughly 20% of sporozoites injected into the host reach the liver stage and ultimately produce merozoites; further emphasising that only a very small percentage of all salivary gland sporozoites establish infection within the host.

However, we note that the numerical reduction in number of sporozoites may be significantly greater than the amount of genetic diversity that is lost, as this is critically dependent on the amount of genetic diversity in the original salivary gland population. Consider that if there are only four unique *P. falciparum* genomes within the salivary glands (which could occur if the vector harboured a single oocyst), then a minimum of only 4 sporozoites is needed to re-establish 100% of genetic diversity within the host. This would be possible even if only tens to hundreds of sporozoites were transmitted. Furthermore, we note that the rate of co-infection in *P. falciparum* has been found to be high: globally it is nearly equivalent to the rate of super-infection in mixed infections carrying two strains [37], and there is evidence that co-infection occurs regularly in high transmission settings [28], observations that would be unlikely if a substantial fraction of diversity was lost with transmission.

As a starting point, we set $p_v$ to 0.2 and $p_h$ to 0.2; though the biological processes impacting the value of $p_h$ differ (it concerns the fraction of gametocytes that are imbibed by a vector), it is subject to similar uncertainties as $p_v$. Studies analysing the genetic diversity present within the vector midgut and salivary glands could strengthen the estimates of these parameters.

**Distribution of number of oocysts ($p_{oocysts}$)**

As described in the main text, we model the number of oocysts that are generated during meiosis in an individual mosquito by a truncated geometric distribution:

$$N_{oocysts} \sim max[Geo(p_{oocysts}), 10]$$

$p_{oocysts}$ therefore represents the probability of having only a single oocyst, and we restrict the maximum number of oocysts to ten. The number of oocysts in wild-caught *Anopheles* mosquitoes can be determined by dissection. We used two references to set $p_{oocysts}$: (i) [5] caught *Anopheles gambiae s.l.* near Kisumu in Kenya, and Fig. 2 of that reference suggests an approximately geometric distribution with $p = 0.5$; (ii) [15] caught *Anopheles gambiae s.l.* near Basse in The Gambiae, and Fig. 4 of that reference appears roughly geometric, with a $p \leq 0.5$. Consequently, we set $p_{oocysts} = 0.5$ in `forward-dream`.

### 3 Supplementary Figures

| Statistic | Definition |
| --- | --- |
| Fraction Mixed Samples | Fraction of samples collected that contain $> 1$ distinct parasites |
| Mean Complexity of Infection (C.O.I.) | Mean number of distinct parasites per sample |
| Number of segregating sites ($S_n$) | Total number of polymorphic sites across all samples |
| Number of singletons | Number of polymorphic sites where the minor allele is present in only one sample |
| Nucleotide diversity ($\pi$) | Mean number of pairwise differences between two samples, normalized to the total number of sites |
| Watterson's Theta ($\theta_w$) | Point estimate of $\theta = 2N_e\mu$ where $\mu$ is the mutation rate, given as $\theta_w = S_n / \sum_{i=1}^{n-1} \frac{1}{i}$ |
| Tajima's $D$ | A test statistic for the neutral mutation hypothesis Tajima1989 |
| Mean IBD | Mean fraction of sites identical between two parasites within the sample |
| Mean IBD track length | Mean length of an IBD track ($\geq 1$ consecutive identical sites) between two parasites within the sample |
| Mean number of IBD tracks | Mean number of IBD tracks between two parasites within the sample |

Table 4: Genetic diversity statistics computed in `forward-dream`.

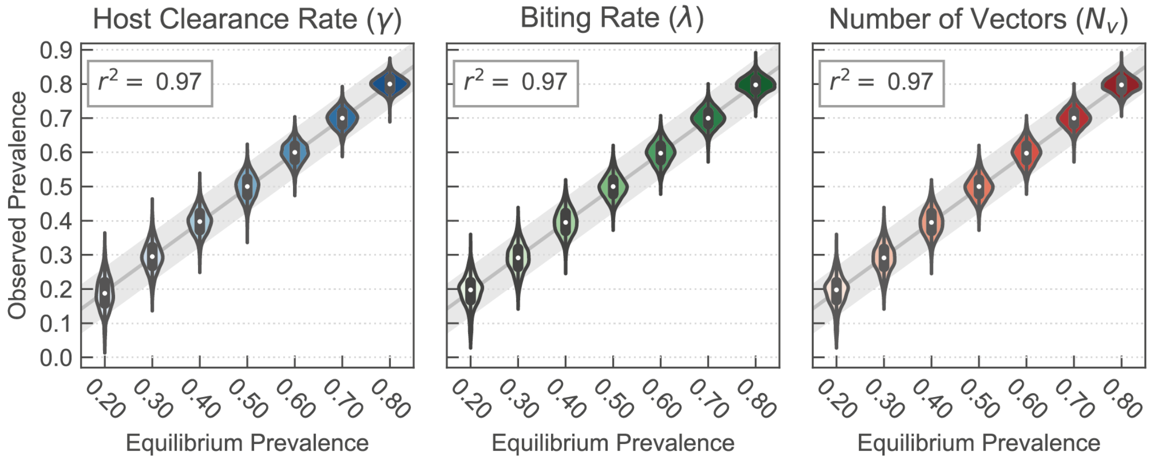


Figure 1: **Validating equilibrium host prevalence values in `forward-dream`.** The epidemiological layer of `forward-dream` implements the Ross-Macondald Model, where the host prevalence is a function of the rate parameters (see Eq. 1 of main text). Violinplots summarize the prevalence values observed in `forward-dream` simulations with expected equilibrium prevalence values varying from 0.2 to 0.8 (computed using Eq. 1) given on the x-axis. The different equilibrium prevalence values were achieved by varying either the host clearance rate ($\gamma$), the vector biting rate ($b$), or the number of vectors ($N_v$). The variance explained ($r^2$) in an ordinary linear regression is shown at top-left of each plot.

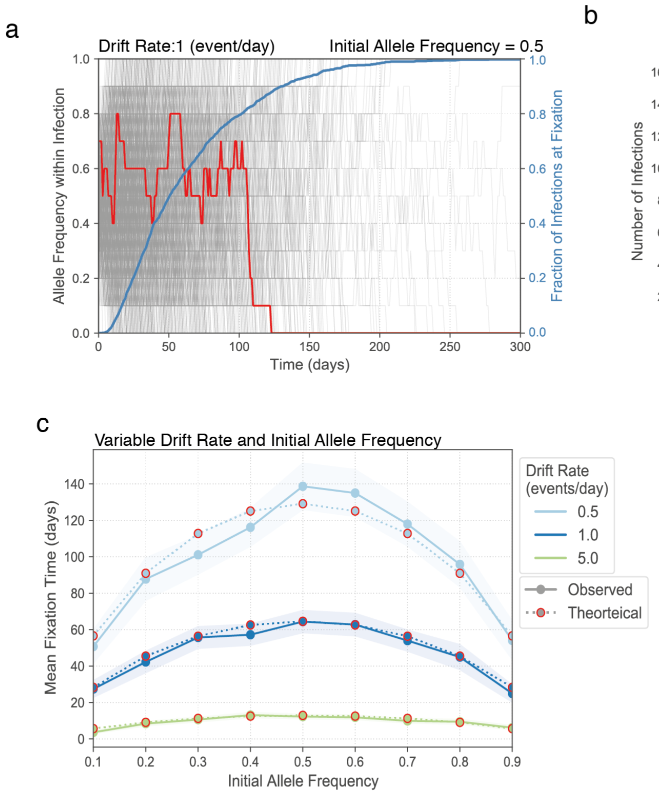


Figure 2: **Validating intra-host fixation times in `forward-dream`.** (a) The infection of a single host is evolved through time and the within-host alelle frequency of a given site is indicated by the red line. The site fixes around day 125. The experiment is repeated 1000 times (grey lines) and the fraction of infections fixed at a given time is indicated by the blue line. All experiments started with an initial allele frequency of 0.5 and a drift rate of 1 event per day. (b) Distribution of fixation times from (a). The observed mean (64.81 days) is very close to the theoretically expected mean from the Moran model (64.56 days) [27]. (c) The experiment in (a) is repeated but with different initial allele frequencies (x-axis) and three different drift rates (light blue, dark blue, and green line). In all cases, the observed mean fixation times are close to the theoretically expected times. Shading gives 95% confidence intervals for mean estimates.

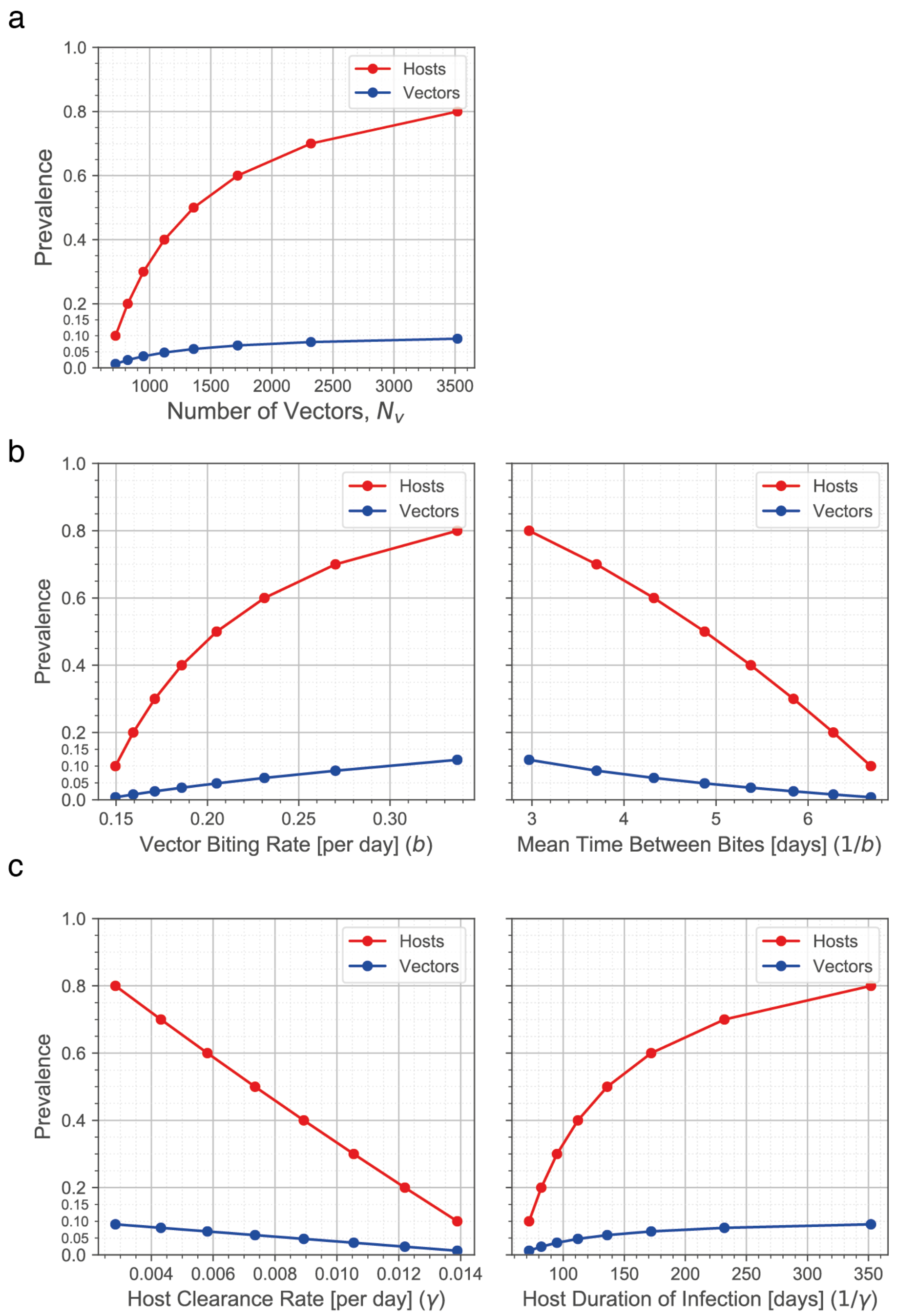


Figure 3: **Varying equilibrium prevalence values in `forward-dream`.** The parameter values of `forward-dream` are varied to produce simulations with equilibrium parasite prevalence ($PfPR$) values varying from 0.2 to 0.8. (a) Varying the number of vectors. Prevalence in hosts indicated in red, vectors in blue. Dots mark parasite prevalence values of 0.1 through 0.8. (b) Varying the vector biting rate $b$. Note $1/b$ gives the average time between successive bites, show in right plot. (c) Varying the host clearance rate ($\gamma$). Note $1/\gamma$ gives the average duration of host infection, shown in right plot.

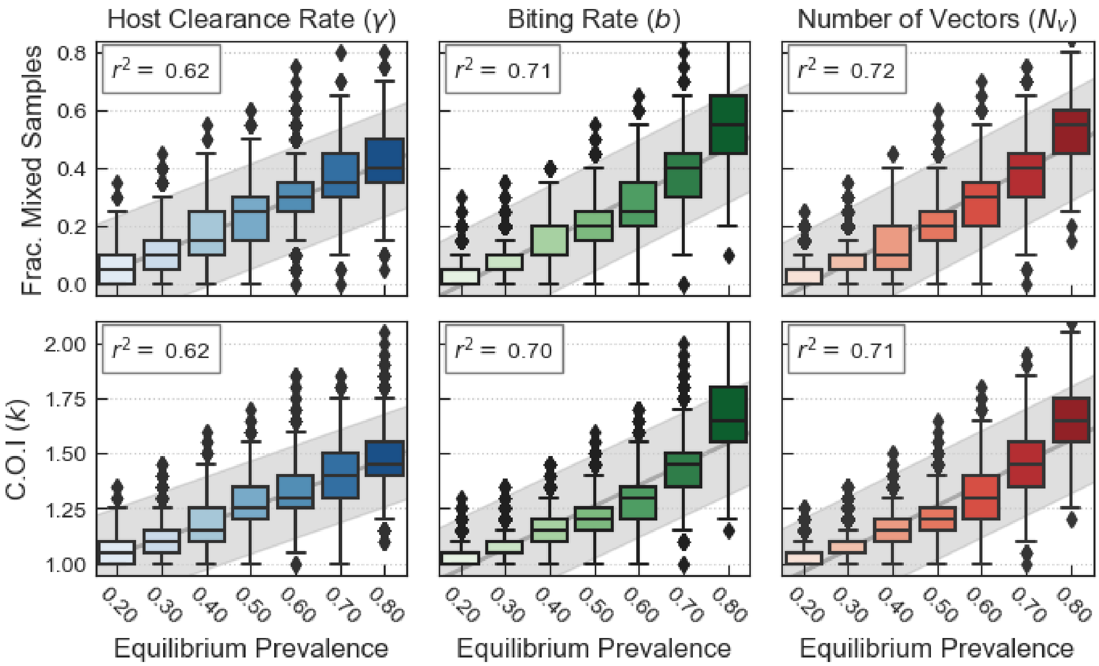


Figure 4: **Equilibrium relationships between parasite prevalence and mixed infection related statistics.** Distributions of mixed infection related genetic diversity statistics (y-axis), plotted for equilibrium parasite prevalence values tuned to between 0.2 and 0.8 (x-axis) in `forward-dream` simulations. Left, middle, and right columns show distributions when parasite prevalence is varied as a function of the host clearance rate ($\gamma$, in blue), vector biting rate ($b$, in green) or number of vectors ($N_v$, in green). Each boxplot contains the result of 30 replicate experiments, where the parasite genomes within 20 randomly selected hosts are collected at every 30 days for 10 years and are used to compute the genetic statistic of interest. The variance explained by ordinary least squares regression is given at top left, and line of best fit and confidence intervals indicated in grey.

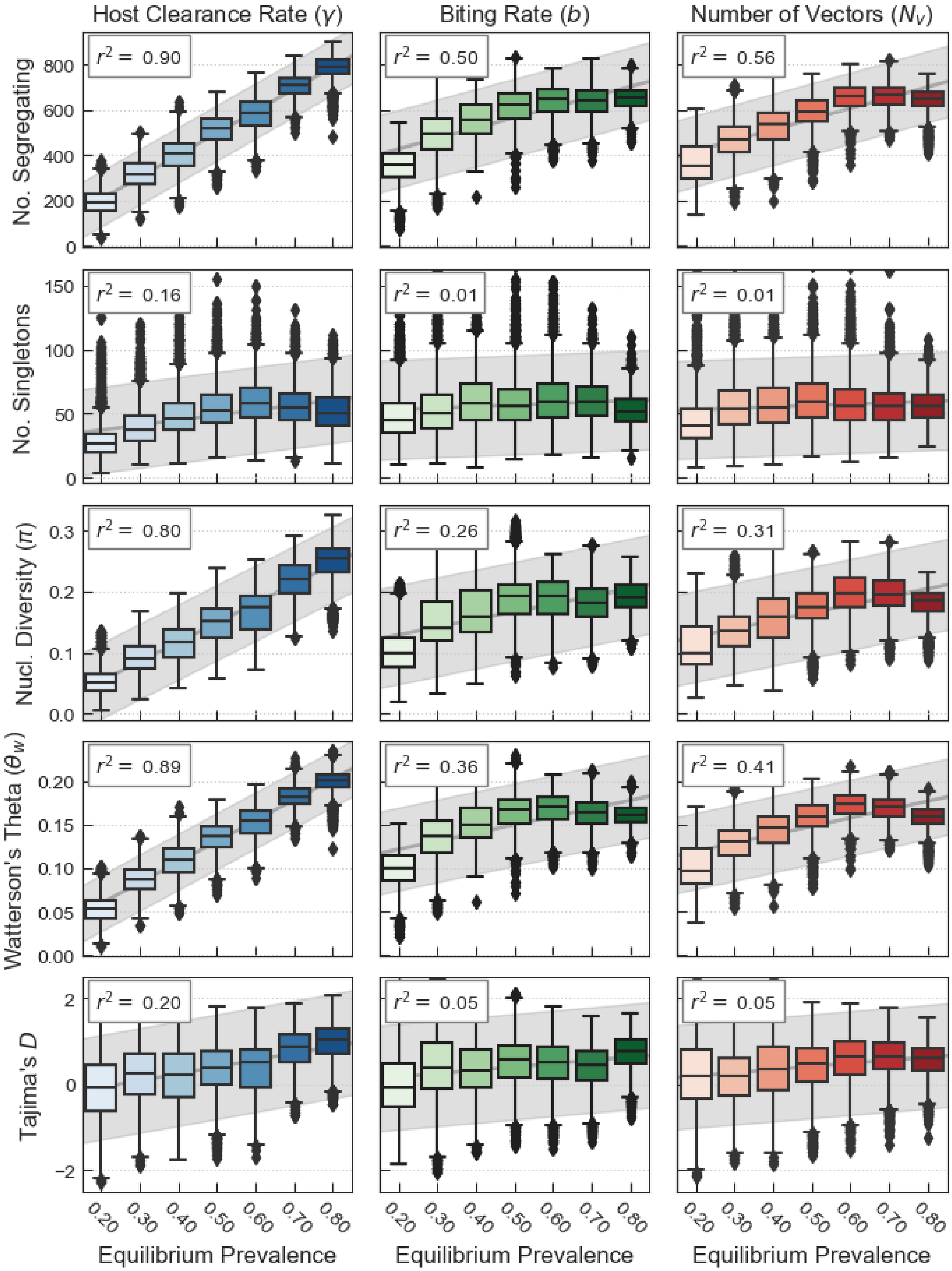


Figure 5: **Equilibrium relationships between parasite prevalence and genetic diversity statistics related to the size and shape of the sample genealogy.** See Figure 4 for details.

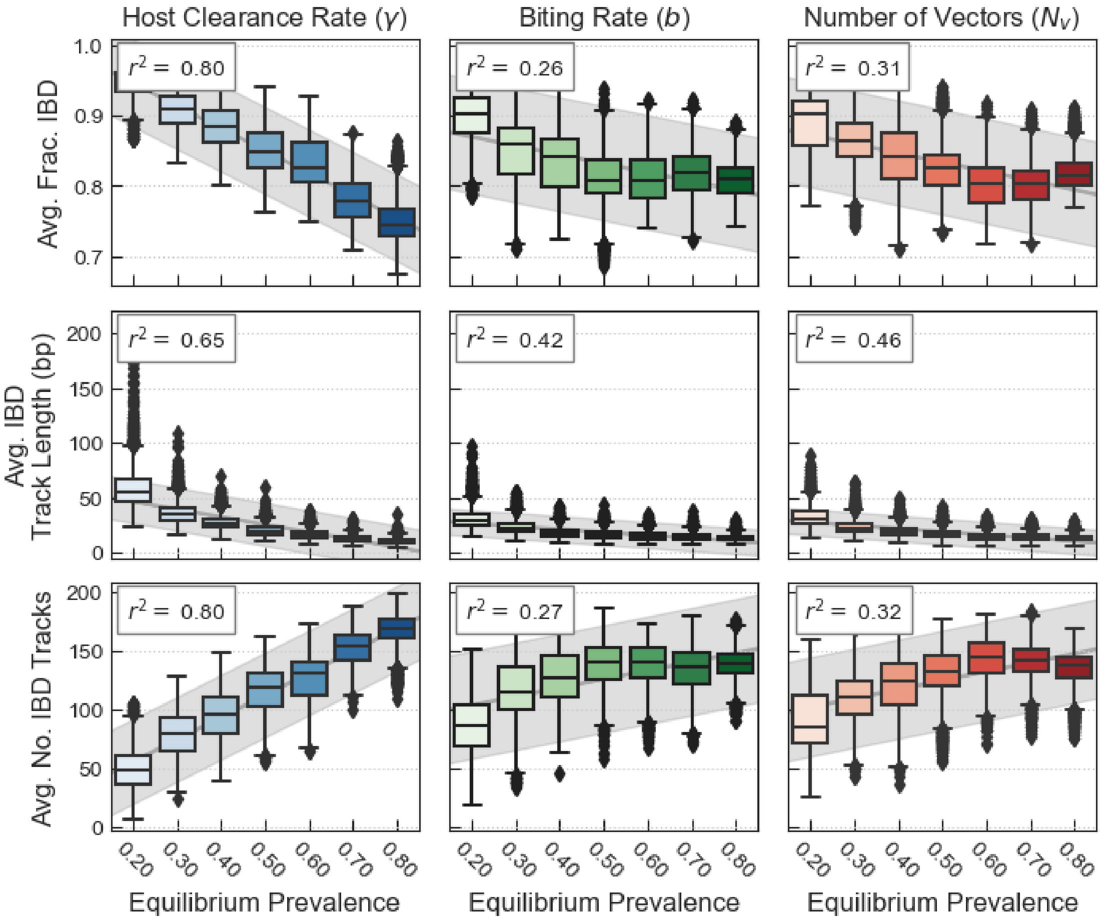


Figure 6: **Equilibrium relationships between parasite prevalence and genetic diversity statistics related to identity-by-descent patterns.** See Figure 4 for details.

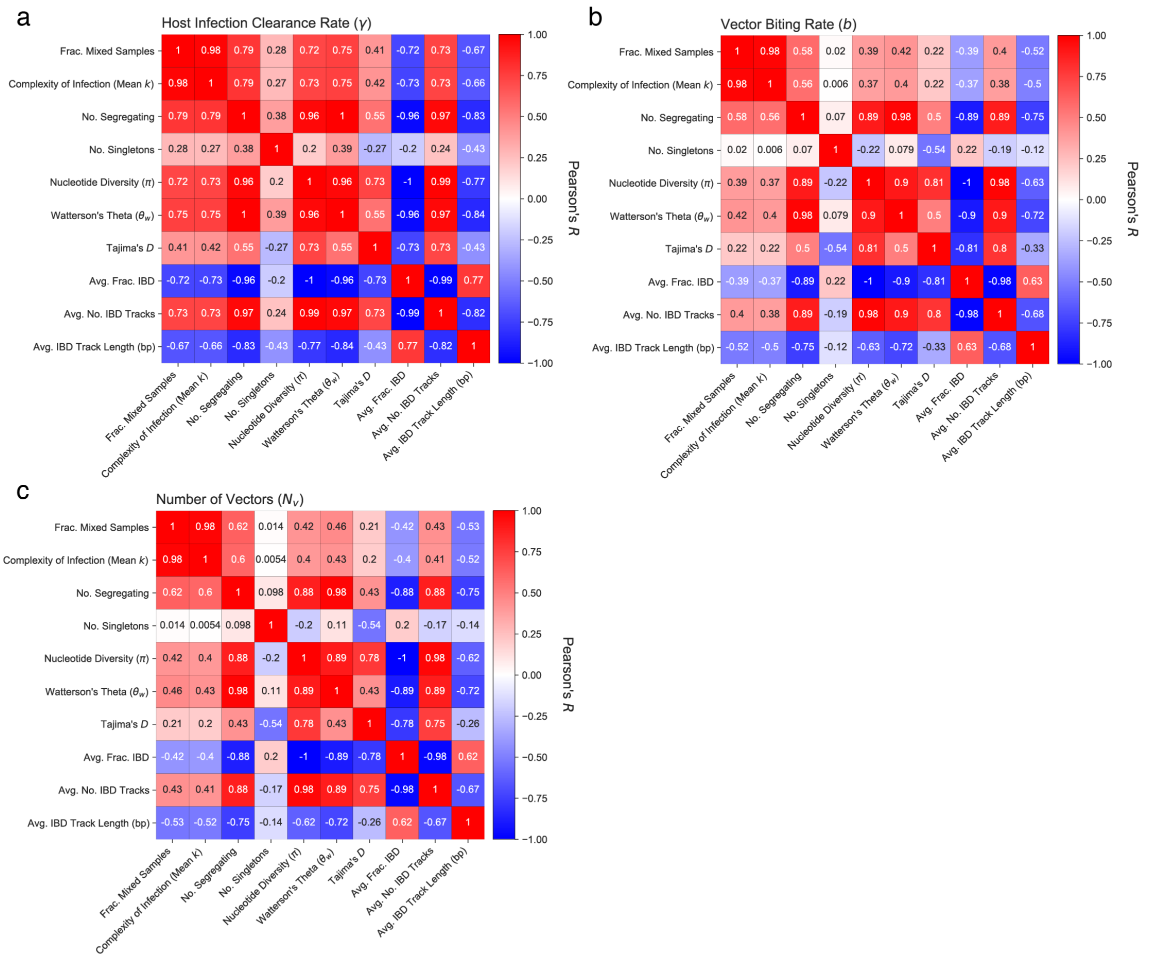


Figure 7: **Co-linearity is observed between different genetic diversity statistics in `forward-dream`** Same data as from Fig 4-Fig **??**. Matrices of Pearson's Correlation Co-efficient ($R$) calculated between all pairs of genetic diversity statistics is shown. In panel (a) host prevalence was tuned to different values between 0.2 and 0.8 by varying the host clearance rate ($\gamma$); (b) by varying the vector biting rate ($b$); or (c) by varying the number of vectors $N_v$. In all cases there is significant co-linearity between different genetic diversity statistics.

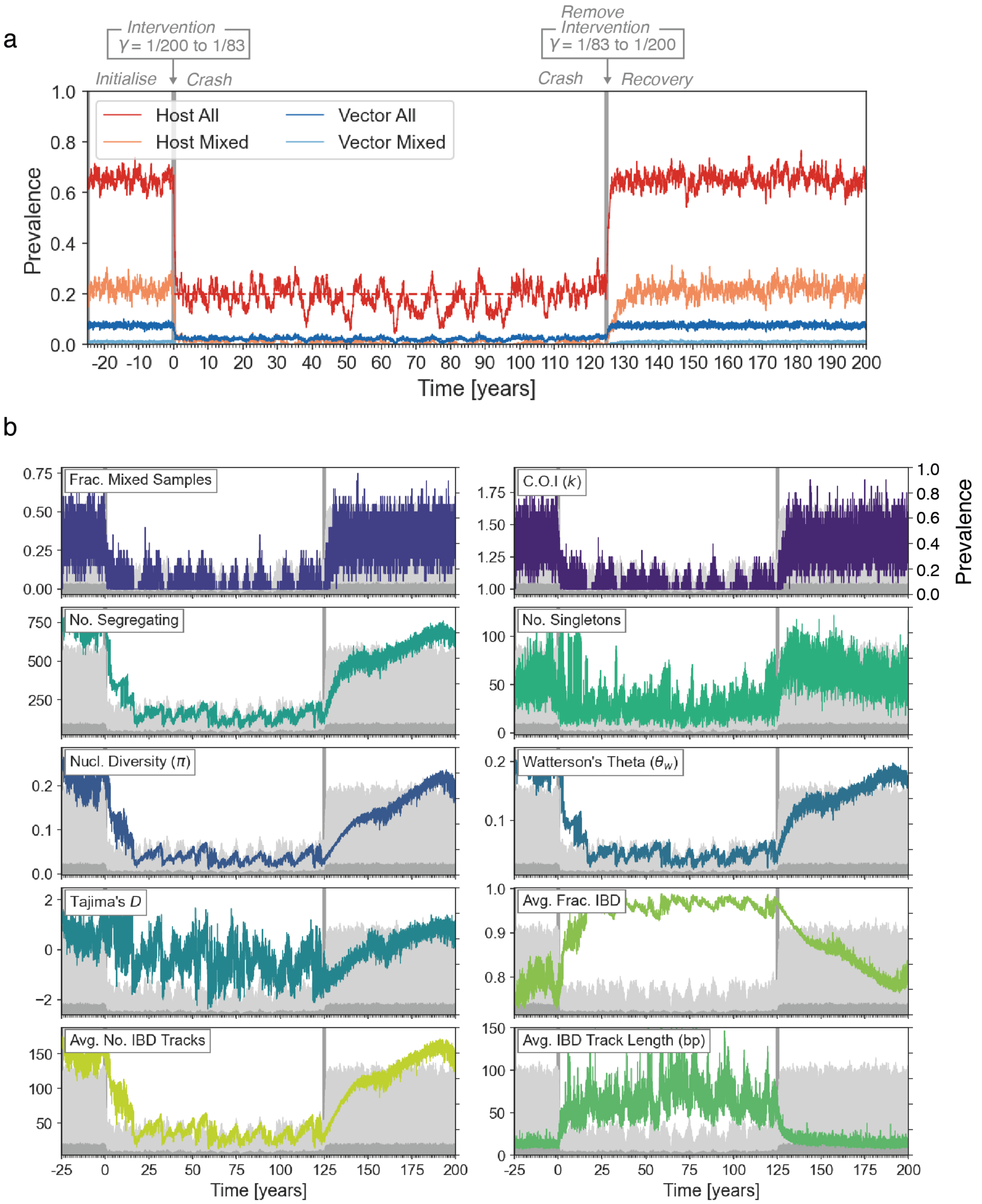


Figure 8: **Behaviour of an individual `forward-dream` simulation during a crash and recovery of parasite prevalence: driven by a change in host clearance rate ($\gamma$).** (a) Prevalence (y-axis) over a 220-year period where a population crash and recovery has occurred. The simulation is first allowed to equilibrate (”Initialise” Epoch). At time zero (x-axis), the host clearance rate is increased from 0.005 (1/200) to 0.012 (1/83), causing a crash in prevalence (”Crash” Epoch). After 125 years (sufficient time to for the simulation to reach a new equilibrium), the host clearance rate is returned to its original value (”Recovery” Epoch). Prevalence of all infected ($k > 0$) and multiply-infected ($k > 1$) hosts is indicated by red and pink lines, respectively. The same is shown for vectors in blues. (b) The same simulatoin as in (a), but with a variety of genetic diversity statistics shown. Note that the statistics are computed from parasite genomes collected from 20 randomly selected hosts every 5 days. For reference, light and dark grey show the host and vector prevalence on the second y-axis.

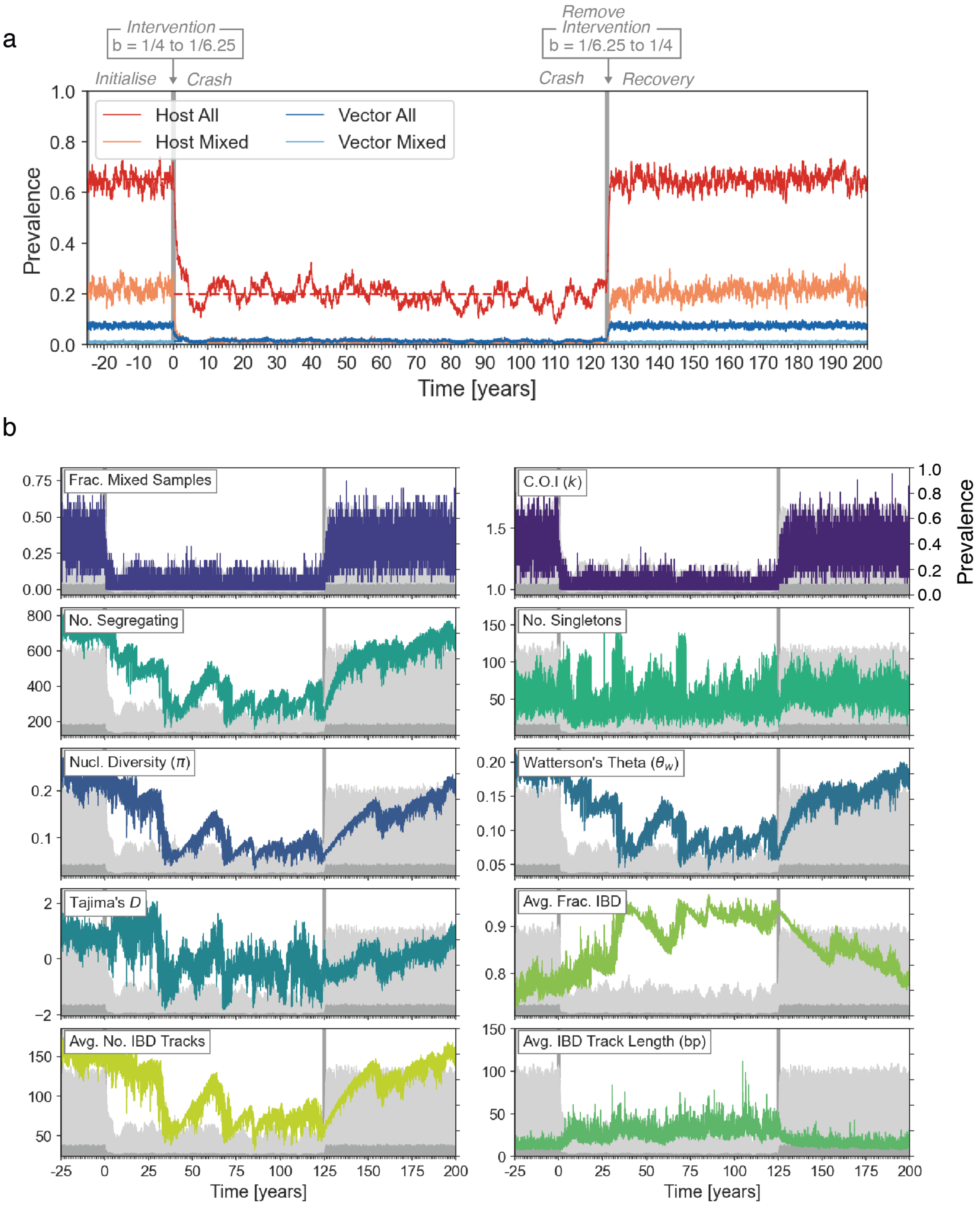


Figure 9: **Behaviour of an individual `forward-dream` simulation during a crash and recovery of parasite prevalence: driven by a change in vector biting rate ($b$).** See Fig 8 for details.

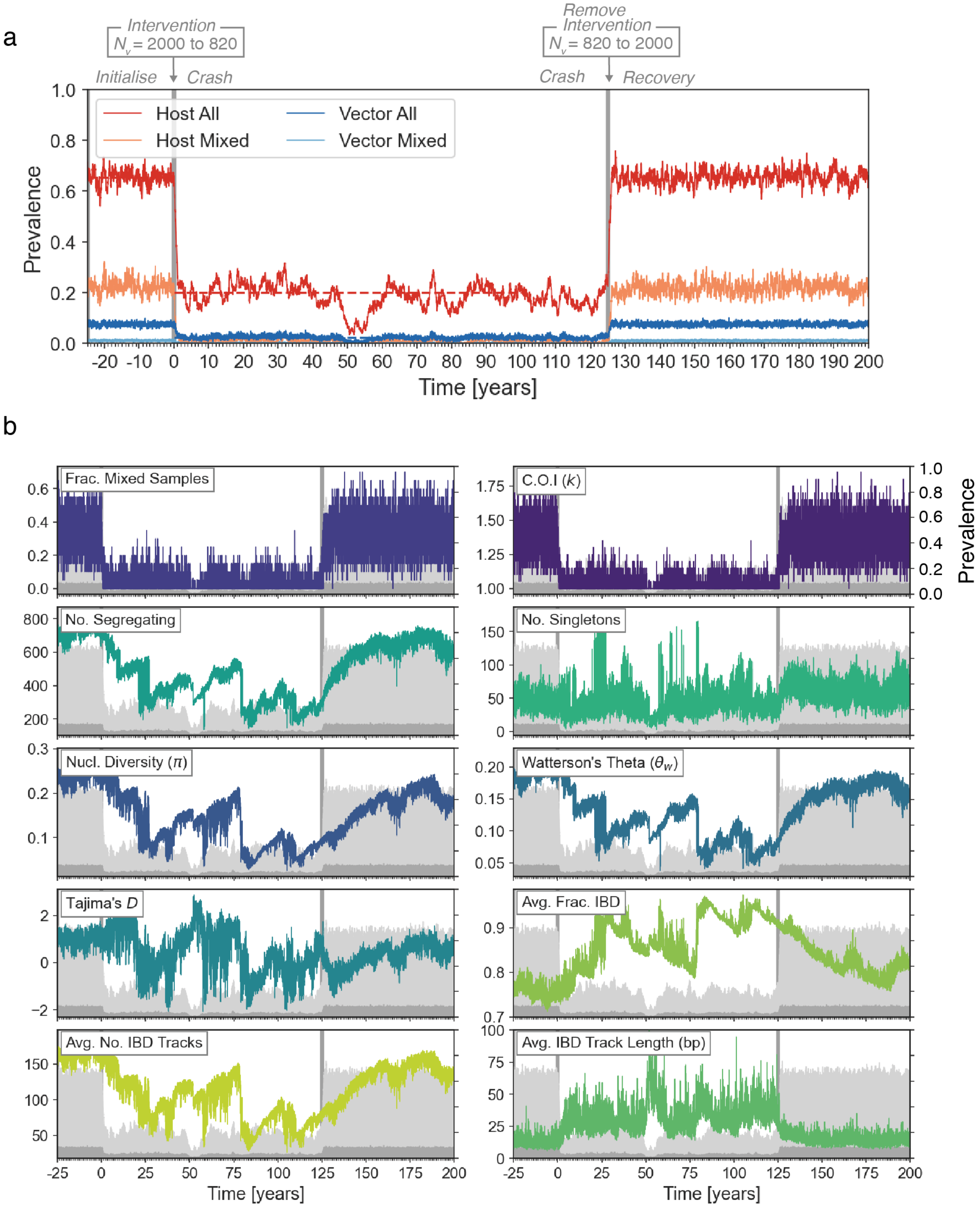


Figure 10: **Behaviour of an individual `forward-dream` simulation during a crash and recovery of parasite prevalence: driven by a change in number of vectors ($N_v$).** See Fig ?? for details.

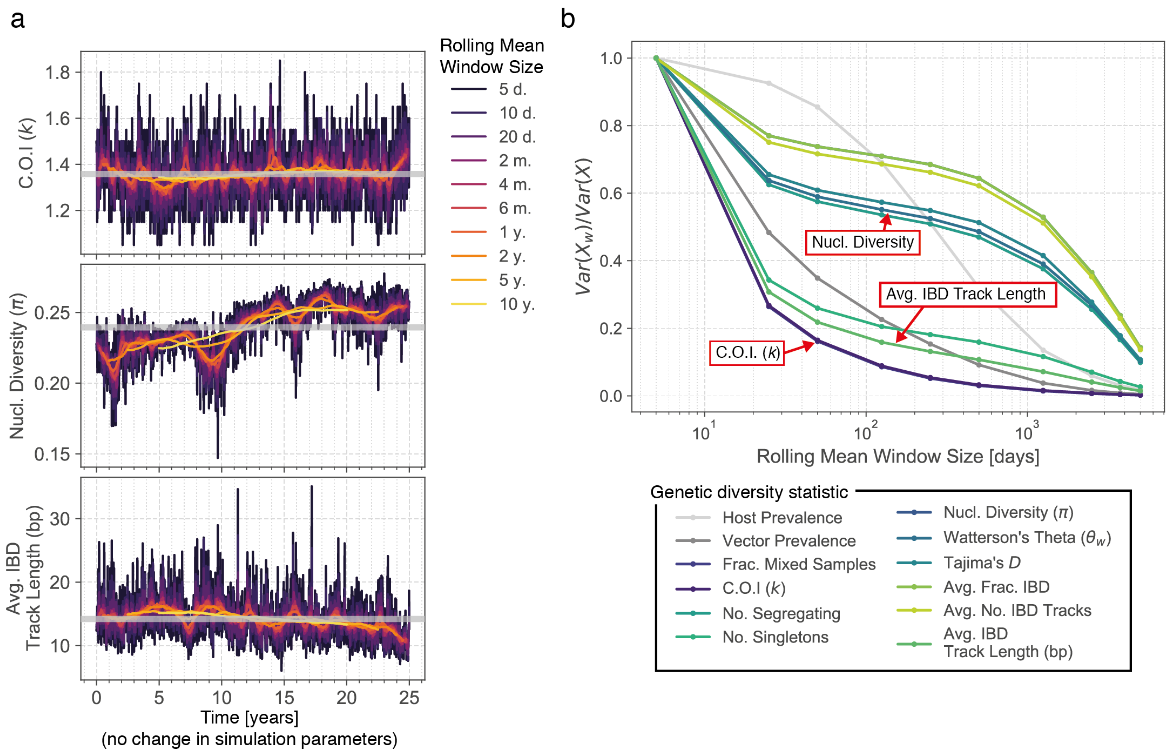


Figure 11: **Temporal fluctuations in genetic diversity statistics of different frequencies, without any change in parasite prevalence.** Panel (a) shows the noisy behaviour of three genetic diversity statistics (y-axis) from an individual simulation where parasite prevalence was kept fixed at 0.65 for a 25 year period (x-axis). The trajectory of each statistic was smoothed using a rolling mean, with window sizes varying from 1 day (1 d., purple), which is equivalent to no smoothing, up to 10 years (10 y., yellow). The mean of the statistic during the 25-year window is indicated with the grey horizontal bar. Notice how even with a 10-year window rolling mean, the nucleotide divesity still deviates from its mean value. (b) Across 100 independent replicate simulations, the reduction in variance of each genetic diversity statistic with increasing rolling mean window sizes is shown. The y-axis gives the ratio of the variance for the window size indicated by the x-axis ($Var(X_w)$) divided by the unsmoothed variance in the genetic diversity statistic ($Var(X)$). Increasing with window size of the rolling mean always reduces the variance, but at different rates for different statistics.

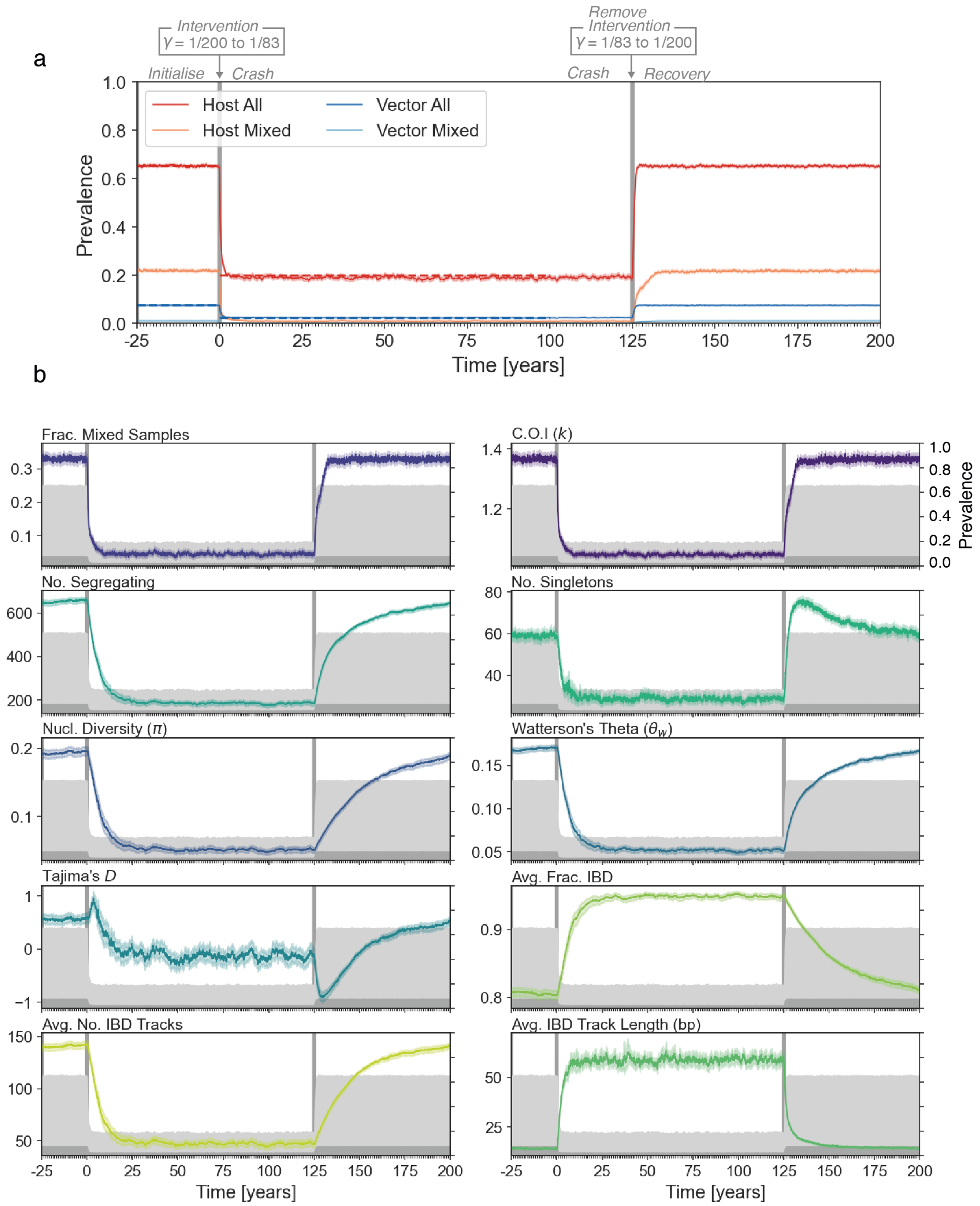


Figure 12: **Average behaviour of 100 replicate `forward-dream` simulations during a crash and recovery of parasite prevalence: driven by a change in host clearance rate ($\gamma$).** Same as Fig 8, but instead of showing an individual simulation trajectory, the averaged trajectory of 100 replicate simulations is shown. Each line is a mean across the 100 replicates, and the shading gives the 95% confidence intervals.

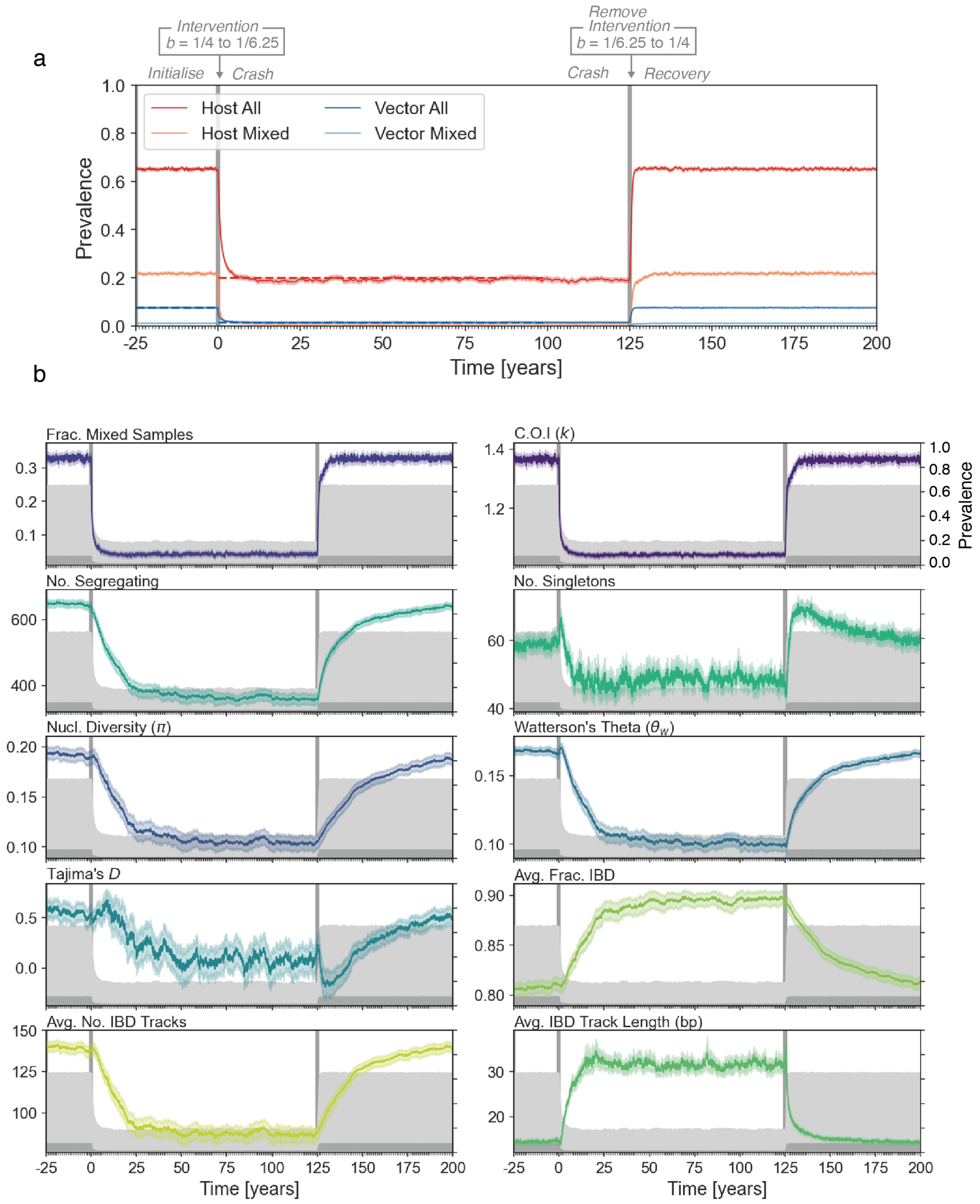


Figure 13: **Average behaviour of 100 replicate `forward-dream` simulations during a crash and recovery of parasite prevalence: driven by a change in vector biting rate ($b$).** Same as Fig 9, but instead of showing an individual simulation trajectory, the averaged trajectory of 100 replicate simulations is shown. Each line is a mean across the 100 replicates, and the shading gives the 95% confidence intervals.

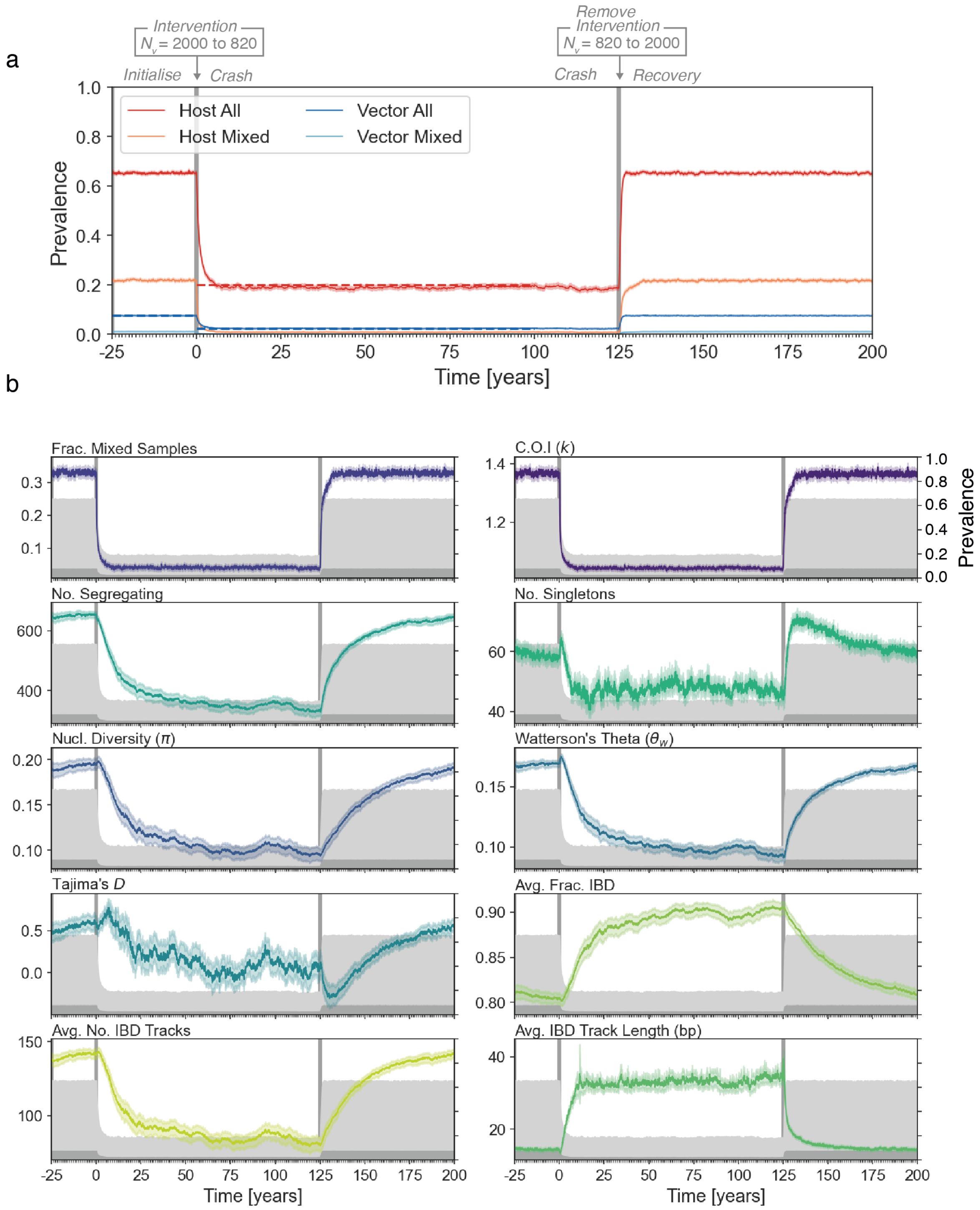


Figure 14: **Average behaviour of 100 replicate `forward-dream` simulations during a crash and recovery of parasite prevalence: driven by a change in the number of vectors ($N_v$).** Same as Fig 10, but instead of showing an individual simulation trajectory, the averaged trajectory of 100 replicate simulations is shown. Each line is a mean across the 100 replicates, and the shading gives the 95% confidence intervals.

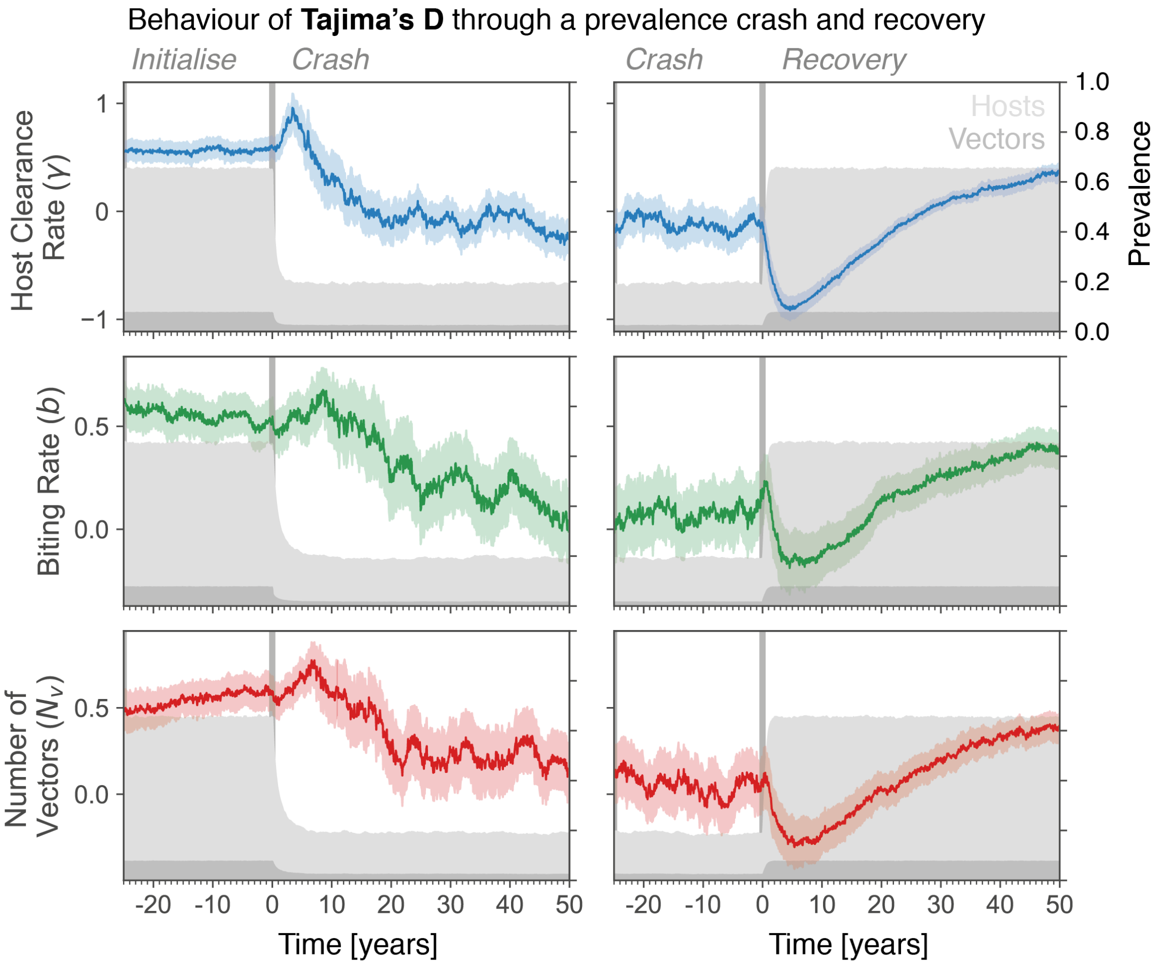


Figure 15: **Average behaviour of Tajima's $D$ during a crash and recovery in parasite prevalence.** Same data as Figures 12-14. Colored lines show mean estimate across 100 replicate simulations, shaded area gives 95% confidence intervals. Notice how Tajima's $D$ increases during a population contraction and decreases during population growth.

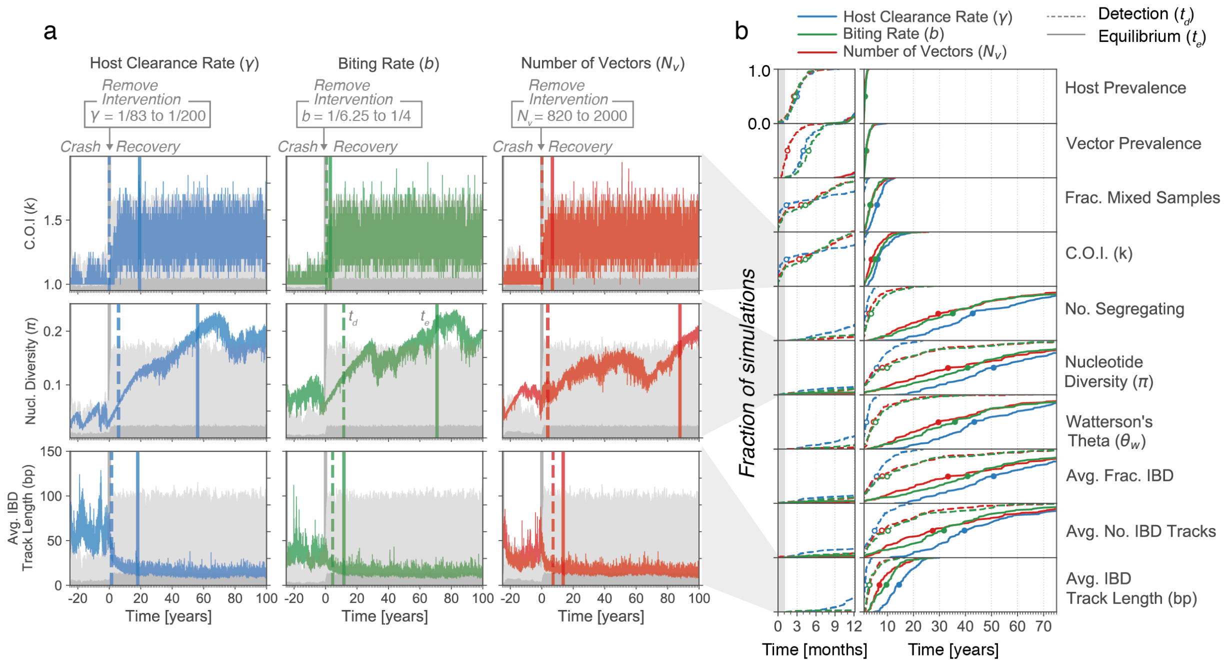


Figure 16: **Detection and equilibrium times of genetic diversity statistics following a recovery in parasite prevalence.** (a) Each plot shows the behaviour of a genetic diversity statistic in an individual simulation through a recovery of parasite prevalence, induced by: left column, reducing the host clearance rate ($\gamma$); middle column, increasing the vector biting rate ($b$); or right column, increasing the number of vectors. The intervention occurs at time zero (x-axis, grey vertical bar) in all cases. For each plot, the detection time (vertical dashed bar) and equilibrium time (vertical solid bar) of the genetic diversity statistic is indicated. Note that here a single simulation is shown for each intervention type. (b) Empirical cumulative density functions (ECDFs) of the detection and equilibrium times of diversity statistics, created from 100 independent replicate simulations for each intervention type. The y-axis gives the fraction of replicate simulations with a detection (dashed line) or equilibrium (solid line) less than the time indicated on the x-axis. Line color specifies the type of intervention. Open and closed circles give medians for the detection and equilibrium times, respectively. The first year is magnified for clarity.

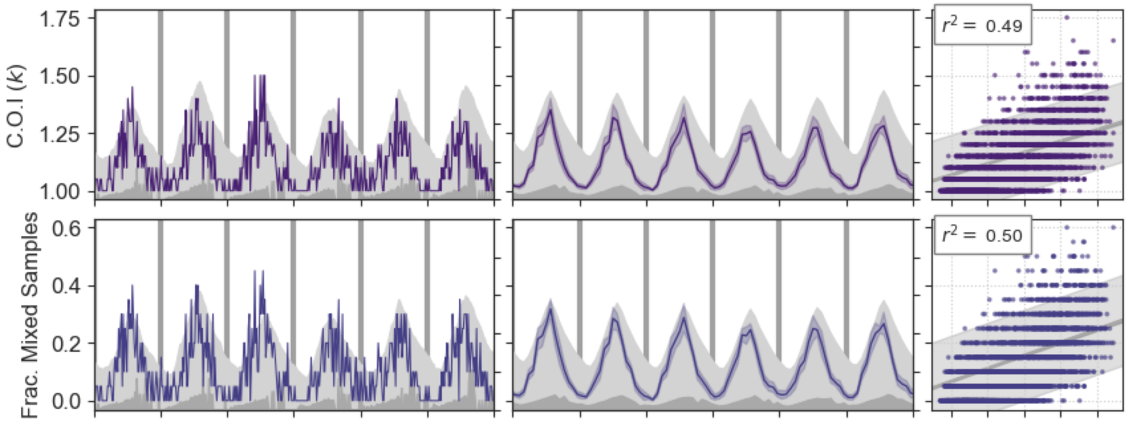


Figure 17: **Responses of mixed infection related genetic diversity statistics to seasonal change in parasite prevalence.** Annual variation in parasite prevalence was induced by varying the number of vectors. The behaviour of genetic diversity statistics for an individual simulation is shown at left. The mean behaviour of 10 independent replicate simulations is shown at middle, with shaded areas giving the 95% confidence intervals. Scatterplots at right show the relationship between each genetic diversity at parasite prevalence across the six years of seasonal fluctuation. Each point represents a genetic diversity estimate (y-axis) computed from sampling parasite genomes from 20 infected hosts in an individual simulation; parasite prevalence (x-axis) is computed across the entire host population at the same time. Data from all 10 replicate simulations has been aggregated. Variance explained $r^2$ from an ordinary linear regression is indicated at top left.

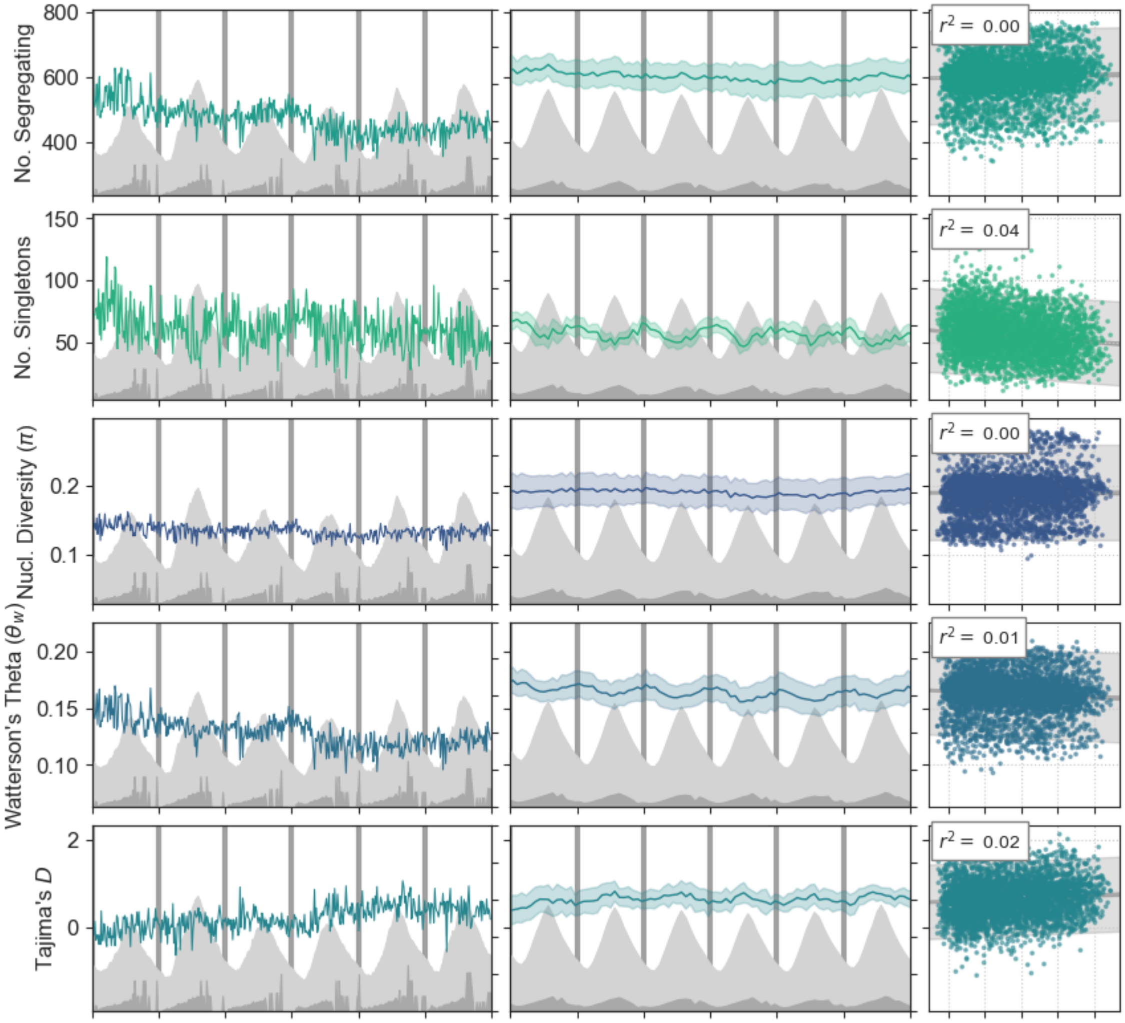


Figure 18: **Responses of sample genealogy related genetic diversity statistics to seasonal change in parasite prevalence.** See Fig 17 for details.

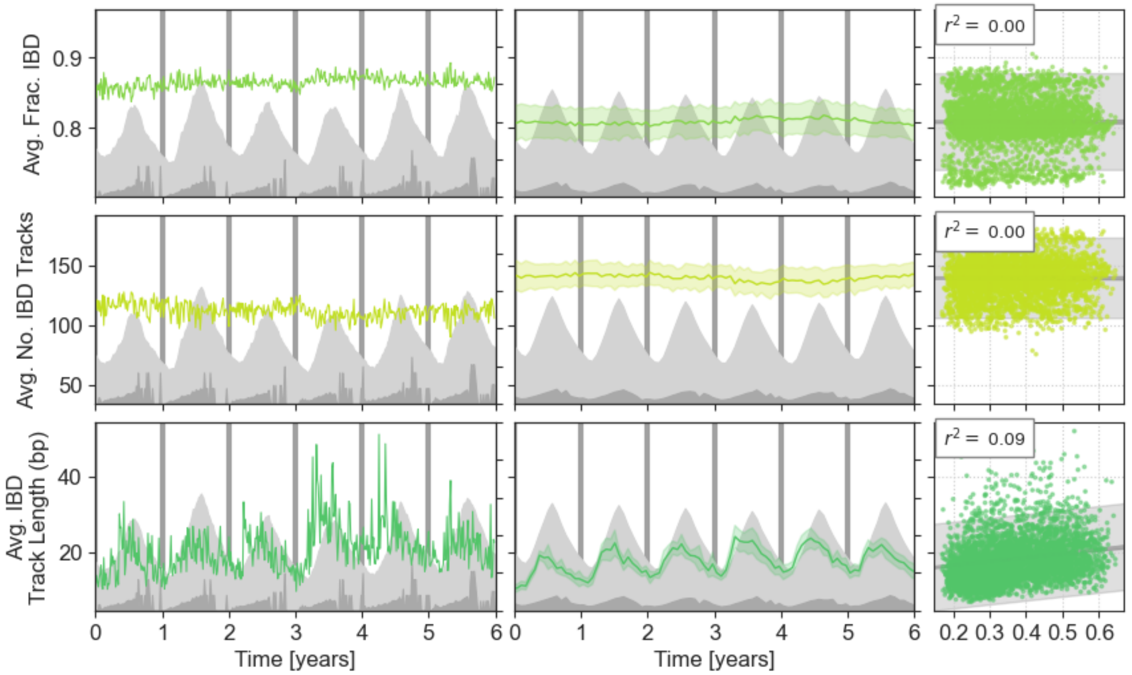


Figure 19: **Responses of IBD related genetic diversity statistics to seasonal change in parasite prevalence.** See Fig 17 for details.

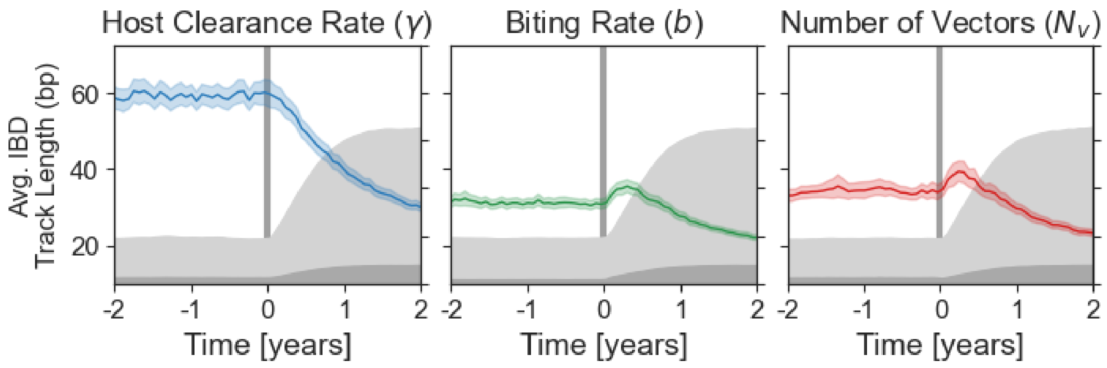


Figure 20: **Zoom on response of average IBD track length to increase in prevalence at beginning of Recovery epoch.** Same data as Fig 12 to Fig 14 but focusing only on the average IBD track length and a four-year window around the beginning of the recovery. Notice how for a change in the number of vectors ($N_v$) there is an increase in average IBD track length at the beginning of the recovery, consistent with epidemic expansion.
